# ZEB1 insufficiency causes corneal endothelial cell state transition and altered cellular processing

**DOI:** 10.1101/547927

**Authors:** Ricardo F. Frausto, Doug D. Chung, Payton M. Boere, Vinay S. Swamy, Huong N.V. Duong, Liyo Kao, Rustam Azimov, Wenlin Zhang, Liam Carrigan, Davey Wong, Marco Morselli, Marina Zakharevich, E. Maryam Hanser, Austin C. Kassels, Ira Kurtz, Matteo Pellegrini, Anthony J. Aldave

## Abstract

The zinc finger e-box binding homeobox 1 (ZEB1) transcription factor is a master regulator of the epithelial to mesenchymal transition (EMT), and of the reverse mesenchymal to epithelial transition (MET) processes. ZEB1 plays an integral role in mediating cell state transitions during cell lineage specification, wound healing and disease. EMT/MET are characterized by distinct changes in molecular and cellular phenotype that are generally context-independent. Posterior polymorphous corneal dystrophy (PPCD), associated with ZEB1 insufficiency, provides a new biological context in which to understand and evaluate the classic EMT/MET paradigm. PPCD is characterized by a cadherin-switch and transition to an epithelial-like transcriptomic and cellular phenotype, which we study in a cell-based model of PPCD generated using CRISPR-Cas9-mediated ZEB1 knockout in corneal endothelial cells (CEnCs). Transcriptomic and functional studies support the hypothesis that CEnC undergo a MET-like transition in PPCD, termed endothelial to epithelial transition (EnET), and lead to the conclusion that EnET may be considered a corollary to the classic EMT/MET paradigm.

## INTRODUCTION

The zinc finger e-box binding homeobox 1 (*ZEB1*) gene encodes a transcription factor involved in epithelial and endothelial cell plasticity critical in development, wound healing and cancer [1]. ZEB1 is a master regulator of cell state transitions (CSTs), namely epithelial to mesenchymal (EMT) or the reverse process, mesenchymal to epithelial (MET). EMT is characterized by distinct molecular and morphologic changes in which epithelial cells lose an epithelial-associated gene expression profile, apicobasal polarity and intercellular adhesions, and gain a mesenchymal-associated gene expression profile and increased migratory capacity. Conversely, the reverse of the EMT process effectively characterizes MET. EMT and MET are tightly regulated CST processes involving the regulation of many genes in a cell-type-independent manner, and for which stable transition states have been identified [2–7]. For example, the cadherin-switch, a well-described feature of EMT, involves the repression of cadherin 1 (*CDH1*; E-cadherin) and activation of cadherin 2 (*CDH2*; N-cadherin) gene expression, with the reverse being observed in MET. In addition, an inverse correlation is observed between the mesenchymal-associated transcription factor ZEB1 and two epithelial-associated transcription factors, ovo-like 2 (OVOL2) and grainy head-like transcription factor 2 (GRHL2), known to directly repress *ZEB1* transcription [6, 8–10].

The corneal endothelium is present on the internal surface of the cornea, which is comprised of three cell types: the external corneal epithelium, the central connective tissue containing a “resting” fibroblast-like cell type (i.e., keratocytes), and the corneal endothelium. The corneal endothelium demonstrates an epithelial organization (i.e., simple squamous epithelium), and expresses both epithelial- and mesenchymal-associated genes [11]. Nevertheless, corneal endothelial cells (CEnC) are considered distinct from most epithelial cell types due to their embryonic origin, unique function and gene expression profile [11]. Therefore, based on anatomic, transcriptomic and functional classification criteria, CEnC may be considered a stable transition cell state between epithelial and mesenchymal cell states. However, this hypothesis remains to be tested, and the classification of CEnC in the context of EMT and MET may be revealed by the important role that ZEB1 plays in the maintenance of the CEnC phenotype.

Posterior polymorphous corneal dystrophy (PPCD) is an autosomal dominant inherited disorder of the corneal endothelium that is characterized by progressive corneal edema and reduced visual acuity. Approximately 30% of affected individuals demonstrate a monoallelic mutation of the *ZEB1* gene, resulting in ZEB1 insufficiency [12]. A smaller percentage of affected individuals demonstrate non-coding mutations in *OVOL2* and *GRHL2*, presumably as a result of ectopic expression of either gene in the corneal endothelium, with subsequent repression of *ZEB1* transcription [13–16]. As a consequence of ZEB1 insufficiency, various epithelial-like features are observed in PPCD corneal endothelium, including a stratified organization, desmosomal intracellular junctions, and expression of an epithelial-like transcriptomic profile, including increased/ectopic expression of epithelial-associated keratins and cadherins (e.g., *CDH1*), and decreased expression of *CDH2* [12, 17, 18]. Recently we reported that reduced ZEB 1 expression in a cell-based model of PPCD using short-interfering RNA (siRNA) targeting ZEB1 resulted in significantly increased CEnC apoptosis and barrier function [18], consistent with prior reports of ZEB1 reduction leading to increased cell death [19, 20] and increased cell barrier function [21–23]. These results provided the first experimental evidence that the corneal endothelium in individuals with PPCD may be characterized by an epithelial-like phenotype not just in form but in function as well. However, given the obvious limitations of using transient siRNA-mediated ZEB1 knockdown to study a condition associated with chronic ZEB1 insufficiency, we generated a constitutive and stable knockdown of ZEB1 protein in an immortalized corneal endothelial cell line utilizing the clustered regularly interspaced short palindromic repeats (CRISPR)-Cas9 gene-editing technology. Herein, we validated the *ZEB1* monoallelic knockout cell line as a cell-based model of PPCD using a transcriptomics approach, and provide evidence (transcriptomic and cell function) to support our hypothesis that a novel MET-like process, termed endothelial to epithelial transition (EnET), best explains the PPCD phenotype. Importantly, key findings from the transcriptomic profiling of human PPCD endothelium [17] were recapitulated in the ZEB1 knockout cell line, further supporting the utility of the CRISPR-Cas9-mediated knockout of ZEB1 in CEnC to gain a better understanding of the molecular factors central to the pathogenesis of PPCD. In addition, based on the evidence provided here for EnET, we propose a corollary to the EMT/MET paradigm, in which EnET is classified as a MET subtype that is characteristic of PPCD.

## RESULTS

### Transcriptomic analysis validates the *ZEB1*^+/-^ CEnC line as a viable cell-based model of PPCD

We developed a *ZEB1*^+/-^ CEnC line to examine the effects of the monoallelic knockout of *ZEB1* on various cellular processes. The mutation introduced by non-homologous end joining repair is a frameshift that generated a premature stop codon, similar to many *ZEB1* mutations associated with PPCD3. Prior to utilizing the CEnC line to study the effects of ZEB1 knockout on cellular processes, we determined the extent to which the line recapitulated one of the primary molecular hallmarks of PPCD endothelium: an ectopic/increased expression of epithelial-specific (and/or associated) genes (Fig 1A). We identified 1715 differentially expressed genes in PPCD endothelium compared to age-matched controls, of which 920 were upregulated and 795 were downregulated [17]. Comparison of the differentially expressed genes in PPCD with genes highly-associated with ex vivo corneal epithelial cells (evCEpC) or with ex vivo corneal endothelial cells (evCEnC) demonstrated that 26% (65/249) of evCEpC genes were upregulated and 37% (40/108) of evCEnC genes were downregulated in PPCD endothelium, significantly different from the expected percentages due to chance alone (p<0.01) (Fig 1A and S1 Fig).

**Fig 1.**
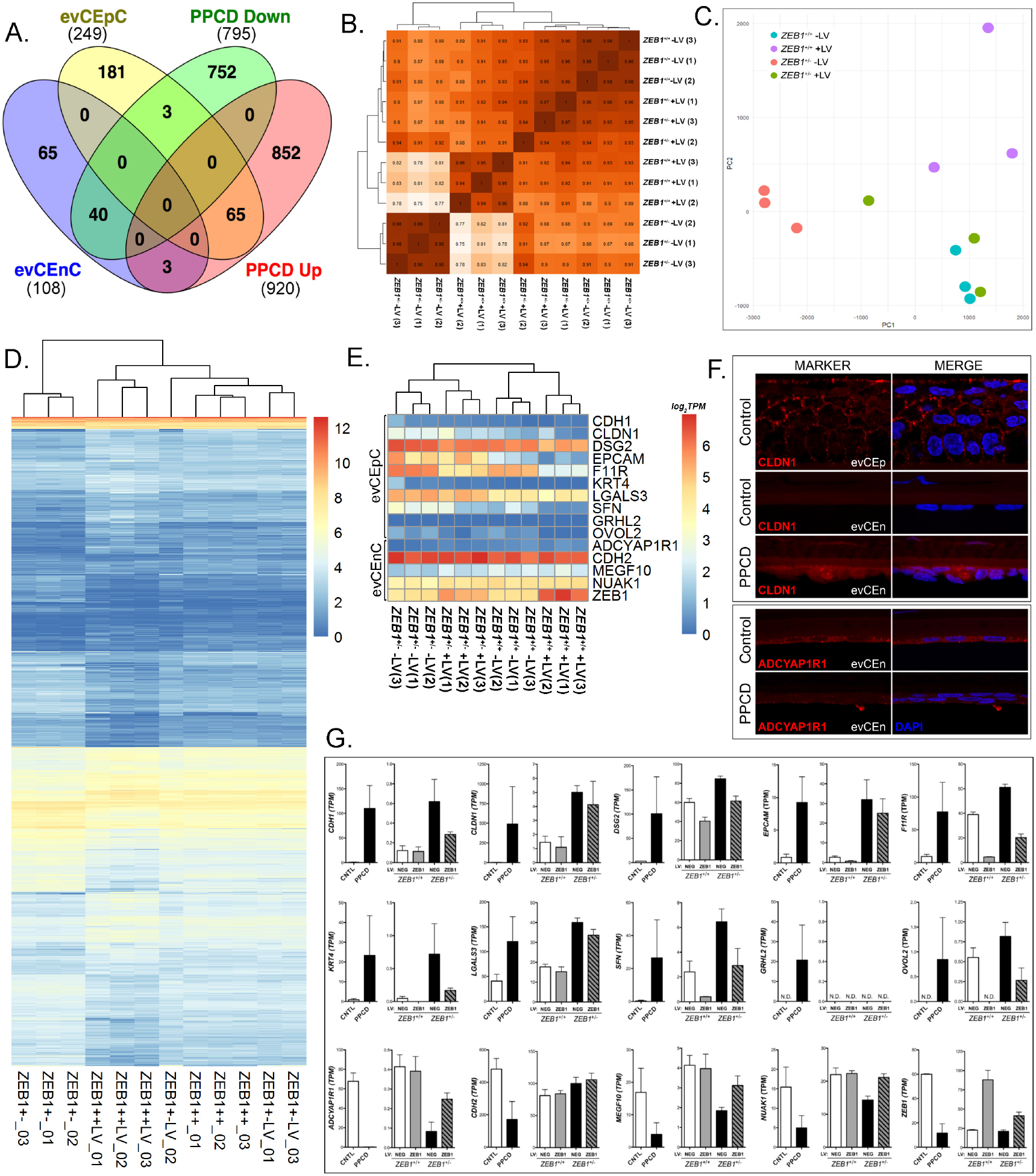
Transcriptomic analysis of the *ZEB1*^+/-^ CEnC line validates it as a model of PPCD. (A) Venn diagram comparing genes specifically expressed by ex vivo corneal epithelial cells (evCEpC) and ex vivo corneal endothelial cells (evCEnC) with differentially expressed genes in PPCD. (B) Spearman correlation heat map, (C) principle component analysis, and (D) heat map demonstrating clustering analysis of the four *ZEB1* CEnC lines and a combined list of 2222 genes that showed significant differential expression in at least one of the three cell lines (*ZEB1*^+/+^ +LV, *ZEB1*^+/-^ -LV, and *ZEB1*^+/-^ +LV) compared with *ZEB1*^+/+^. (E) Hierarchical clustering heat map of selected epithelial- and endothelial-specific genes and *ZEB1* CEnC lines. (F) Immunofluorescence showing expression of the epithelial-associated protein CLDN1 and the corneal endothelial-associated protein ADCYAP1R1 in PPCD endothelium. Expression of CLDN1 in corneal epithelium (evCEp) was used as a positive control. CLDN1 and ADCYAP1R1 were visualized with Alexafluor 594 (red), and nuclei were stained with DAPI (blue). (G) Bar graphs showing the expression of selected epithelial- and endothelial-specific genes (see (E)) in PPCD endothelium and in the *ZEB1* CEnC lines. Gene expression is given in TPMs.

To study the effects of reconstitution of ZEB1 expression on the corneal endothelial transcriptome, we generated stable transgenic *ZEB1*^+/+^ and *ZEB1*^+/-^ cell lines expressing exogenous ZEB1 by the introduction of *ZEB1* cDNA using lentivirus containing the transgene. Three independent cell clones for each of the four cell lines (*ZEB1*^+/+^ -LV, *ZEB1*^+/+^ +LV, *ZEB1*^+/-^ -LV and *ZEB1*^+/-^ +LV) were generated for a total of 12 samples. To examine the relationship of the four *ZEB1* CEnC lines to each other we compared a list of 2222 differentially expressed genes (defined by differential expression in one or more of three cell lines (*ZEB1*^+/+^ +LV, *ZEB1*^+/-^ -LV and *ZEB1*^+/-^ +LV) compared to the reference cell line (*ZEB1*^+/+^ -LV)) to all genes expressed in the 12 samples by Spearman correlation (Fig 1B). The *ZEB1*^+/+^ -LV and the *ZEB1*^+/-^ -LV groups showed a correlation of ~0.89. Reconstitution of the *ZEB1*^+/-^ cell line with ZEB1 (*ZEB1*^+/-^ +LV) increased its correlation with *ZEB1*^+/+^ -LV to ~0.95, the highest correlation between any two groups. The lowest correlation was demonstrated between the *ZEB1*^+/-^ -LV and *ZEB1*^+/+^ +LV cell lines, where the difference in ZEB1 abundance is the greatest, with a correlation of ~0.79. Principle component analysis was also used to assess the relationship of the samples to each other based on the expression of the 2222 genes defined as differentially expressed (Fig 1C). In general, the three samples from each group clustered with each other. While distinct clusters for the *ZEB1*^+/+^ -LV, *ZEB1*^+/-^ -LV and *ZEB1*^+/+^ +LV were observed, the *ZEB1*^+/-^ +LV were clustered more closely to the to the *ZEB1*^+/+^ -LV cells than the other two groups. Hierarchical clustering and heatmap of the 12 samples demonstrated similar results to those observed with Spearman correlation and PCA, where the *ZEB1*^+/+^ -LV and *ZEB1*^+/-^ -LV groups demonstrate distinct clusters, with the *ZEB1*^+/-^ +LV group having a stronger association with the *ZEB1*^+/+^ -LV group (Fig 1D).

To determine if the *ZEB1*^+/-^ CEnC line sufficiently recapitulates the epithelial-like gene expression observed in PPCD3, we compared the expression of a random selection of corneal epithelial-(CDH1, *CLDN1, DSG2, EPCAM, F11R, KRT4, LGALS3, SFN, GRHL2* and *OVOL2*) and endothelial-(*ADCYAP1R1, CDH2, MEGF10, NUAK1, ZEB1*) associated genes that are differentially expressed in PPCD across the four CEnC lines (Fig 1E). Hierarchical clustering analysis of the four *ZEB1* CEnC groups against the 15 selected genes demonstrated that the samples within each group clustered together, but the two main branches clustered based on the *ZEB1* genotype (*ZEB1*^+/+^ or *ZEB1*^+/-^). Although the differential expression of the selected corneal epithelial- or endothelial-associated genes in PPCD was previously identified (Fig 1A), the expression of the encoded protein has not been determined for a majority of these genes. Due to the scarcity of corneal endothelial tissue from affected individuals, we assessed protein expression of one epithelial-associated protein, claudin 1 (CLDN1), and of one endothelial-associated protein, adenylate cyclase activating polypeptide 1 receptor type 1 (ADCYAP1R1), in PPCD endothelium (Fig 1F). While ectopic CLDN1 expression was observed in PPCD endothelium, ADCYAP1R1 expression was markedly decreased in PPCD endothelium compared with normal endothelium. Ninety percent (9/10) of the selected corneal epithelial-associated genes showing increased/ectopic expression in PPCD endothelium also demonstrated increased/ectopic expression in *ZEB1*^+/-^ -LV cells; *GRHL2* was not expressed (Fig 1G). Similarly, eighty percent (4/5) of the corneal endothelial-associated genes showing decreased expression in PPCD endothelium also demonstrated decreased expression in *ZEB1*^+/-^ -LV cells; CDH2 expression was increased.

### ZEB1 insufficiency induces morphologic changes in cultured CEnC

The effects of ZEB1 insufficiency on cell morphology were analyzed using phase-contrast microscopy (Fig 2). Cells for each cloned line were examined at sub-confluent (Fig 2A) and confluent densities (Fig 2B). Most of the sub-confluent *ZEB1*^+/+^ -LV cells demonstrated a cobblestone-like morphology with a few cells demonstrating bipolar morphology. In contrast, the sub-confluent *ZEB1*^+/-^ -LV cells demonstrated a polygonal/cobblestone-like morphology, no bipolar morphology and grew in discrete patches. The sub-confluent *ZEB1*^+/-^ +LV cells, which were reconstituted with ZEB1, demonstrated morphologic characteristics similar to the *ZEB1*^+/+^ - LV cells.

**Fig 2.**
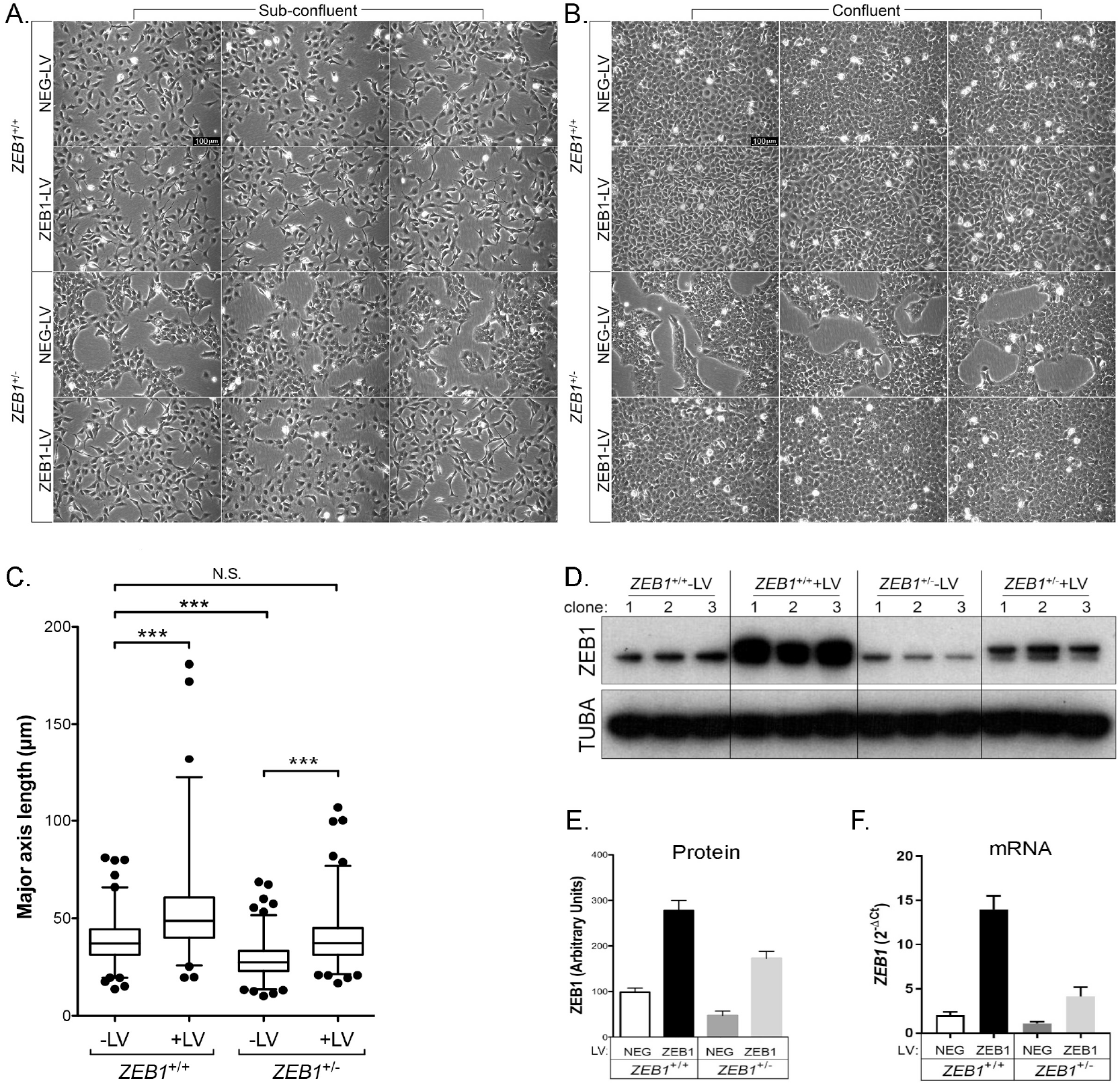
ZEB1 regulates CEnC morphology in a manner consistent with EMT. (A) Sub-confluent cultures established 1 day after seeding of the two control and two *ZEB1* transgenic CEnC lines with each genotype (*ZEB1*^+/+^ -LV, *ZEB1*^+/+^ +LV, *ZEB1*^+/-^ -LV and *ZEB1*^+/-^ +LV) represented by three independent clones (individual images). (B) Confluent cultures of cell line clones shown in (A) established 3 days post-seeding. Scale bars in (A) and (B) represent 100μm distance. (C) Box and whiskers plot showing the cell major axis length (MAL) distribution for each of the CEnC lines. MAL was used to assess cell morphology as a measure of cell state phenotype. Note that a relatively short MAL is indicative of an epithelial-like phenotype, while a relatively long MAL is indicative of a mesenchymal-like phenotype. Box encompasses 50% of data points, line in box is the median of the MAL and whiskers encompass 98% of data points (n=357-688). Comparisons of the MAL for the CEnC lines were performed using one-way ANOVA with post-hoc Tukey test. ***, P<0.001; N.S., not significant (p>0.05). (D) Western blot showing ZEB1 levels in the twelve clones (3 independent clones per genotype) used in this study. Alpha-tubulin (TUBA) was used as a loading control. (E) ZEB1 protein abundance was determined by densitometric analysis of Western blot data shown in (D). Quantification data are represented as mean ±SEM (n=3). Bar graphs showing ZEB1 protein (E) and *ZEB1* mRNA (F) abundances in the four CEnC lines. ZEB1 transcript abundance was measured relative to GAPDH and plotted as 2^-ΔCt^.

Analysis of the transgenic cells at a time point where the clones from 3 of 4 cell groups established a confluent monolayer revealed morphologic changes in the *ZEB1*^+/+^ +LV and *ZEB1*^+/-^ -LV cells compared with the *ZEB1*^+/+^ -LV cells (Fig 2B). Similar to the normal CEnC monolayer on the posterior surface of the cornea, CEnC in 2D culture form a monolayer of polygonal shaped cells, a characteristic observed in *ZEB1*^+/+^ -LV cells. The *ZEB1*^+/-^ -LV cells did not form a contiguous monolayer, but instead maintained distinct patches (albeit covering a larger area) of cell growth and robust cobblestone-like morphology. Reconstitution of the *ZEB1*^+/-^ cells with ZEB1 (*ZEB1*^+/-^ +LV) resulted in the formation of a contiguous monolayer reminiscent of the *ZEB1*^+/+^ -LV confluent cells, thus re-establishing an endothelial-like phenotype without propelling them to a fibroblast-like phenotype, which occurred in *ZEB1*^+/+^ cells in which ZEB1 levels were augmented (i.e., *ZEB1*^+/+^ +LV).

To assess for significant differences in the morphologic characteristics of the cells in each group, we measured the major-axis length (MAL) of each cell in the sub-confluent cultures for the three clones in each group and graphed the data as a box-plot (Fig 2C). We used the MAL of a cell as an indirect measure of the cell state within the epithelial to fibroblastic spectrum of cell states, so that a short MAL was characteristic of epithelial cell morphology and long MAL was characteristic of fibroblast cell morphology. *ZEB1*^+/-^ -LV cells had a mean MAL of 28.6 μm (range: 10.1-68.7 μm), significantly less than the mean MAL for *ZEB1*^+/+^ -LV (38.5 μm; range: 13.7-81.1 μm) (p<0.001). Reconstitution of ZEB1 in the *ZEB1*^+/-^ cells (*ZEB1*^+/-^ +LV cells) resulted in a mean MAL of 39.4 μm (range: 16.8-106.9 μm), significantly increased compared with the mean MAL for *ZEB1*^+/-^ -LV cells (p<0.001), and not significantly different compared with the *ZEB1*^+/+^ -LV cells (p>0.05). The mean MAL increased further to 53.0 μm (range: 19.6-180.8 μm) in the *ZEB1*^+/+^ +LV cells, significantly increased compared with the *ZEB1*^+/+^ -LV cells (p<0.001). Collectively, all of the observed morphologic changes were directly correlated with ZEB1 protein (Fig 2D and 2E) and *ZEB1* mRNA (Fig 2F) levels.

### Reduced ZEB1 levels lead to decreased CEnC migration capacity

A non-wounding cell migration assay was performed to assess the effect of reduced ZEB1 levels on CEnC migration capacity (Fig 3). Phase-contrast microscopy demonstrated that reduction of ZEB1 (*ZEB1*^+/-^ -LV) markedly reduced CEnC migration capacity (~24% gap closure) compared with control CEnC (*ZEB1*^+/+^ -LV; ~83% gap closure; p<0.001). Reconstitution of *ZEB1*^+/-^ cells with ZEB1 (*ZEB1*^+/-^ +LV) appeared to rescue the attenuated migratory phenotype observed in *ZEB1*^+/-^ -LV CEnC (p<0.001), with a gap closure (~74%) that was not significantly different from that in the *ZEB1*^+/+^ -LV CEnCs (p>0.05). In contrast, augmentation of ZEB1 levels in *ZEB1*^+/+^ cells (ZEB1+++LV) did not result in a significant increase in cell migration capacity (~84% gap closure) compared with *ZEB1*^+/+^ -LV cells (~83% gap closure) (p>0.05).

**Fig 3.**
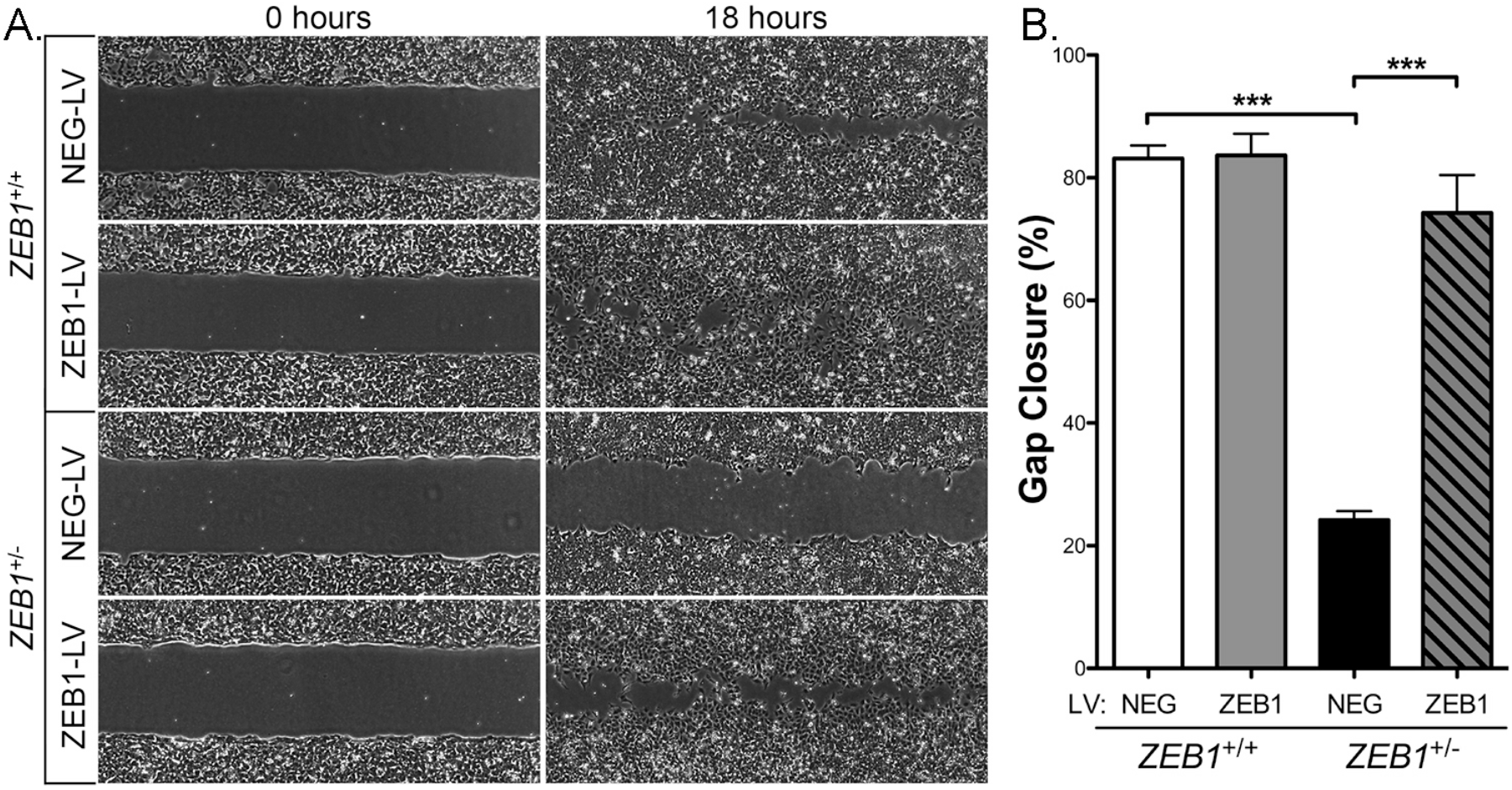
ZEB1 reduction impairs CEnC migration capacity. (A) Representative images at 0 hours showing a gap of ~500μm (width) and at 18 hours showing degree of cell gap closure (i.e., cell migration) for each of the control (*ZEB1*^+/+^ NEG-LV and *ZEB1*^+/-^ NEG-LV) and *ZEB1* transgenic (*ZEB1*^+/+^ ZEB1-LV and *ZEB1*^+/-^ ZEB1-LV) CEnC lines. (B) Bar graph showing percent of gap closure at 18 hours. Data are represented as the mean ±SEM (n=12). Comparisons were performed using one-way ANOVA with post-hoc Tukey test. ***, P<0.001.

### Reduced ZEB1 levels leads to decreased CEnC proliferation capacity

To determine the effect that decreased ZEB1 has on CEnC proliferation, we measured cell proliferation in *ZEB1*^+/+^ and *ZEB1*^+/-^ cell lines with transient ZEB1 lentivirus transduction (Fig 4). Two days after transduction, some of the cells were re-seeded to assess cell proliferation while the remaining cells were lysed and prepared for Western blotting. ZEB1 Western blot confirmed the expected relative ZEB1 protein levels in each of the four groups (Fig 4A). Cells from the newly seeded cultures were collected at 3, 48, 72 and 96 hours and counted. A ratio (N_t_/N_0_) of cell number at time t (N_t_ = 48, 72 or 96 hours) versus the cell number at 3 hours (defined as the reference, N_0_) was graphed as a measure of cell proliferation (Fig 4B). At 72 and 96 hours, the *ZEB1*^+/-^ -LV cells demonstrated significantly less cell proliferation compared with the *ZEB1*^+/+^ - LV cells (p<0.0001). Reconstitution of *ZEB1*^+/-^ cells with ZEB1 (*ZEB1*^+/-^ +LV) resulted in a significant increase in cell proliferation at 72 hours (p<0.5) and 96 hours (p<0.0001) compared with *ZEB1*^+/-^ -LV, returning to a level that was not significantly different from that of the *ZEB1*^+/+^ -LV cells (p>0.05). Consistent with the above results, the *ZEB1*^+/+^ +LV cells demonstrated a significant increase in proliferation compared with the *ZEB1*^+/+^ -LV cells (p<0.0001).

**Fig 4.**
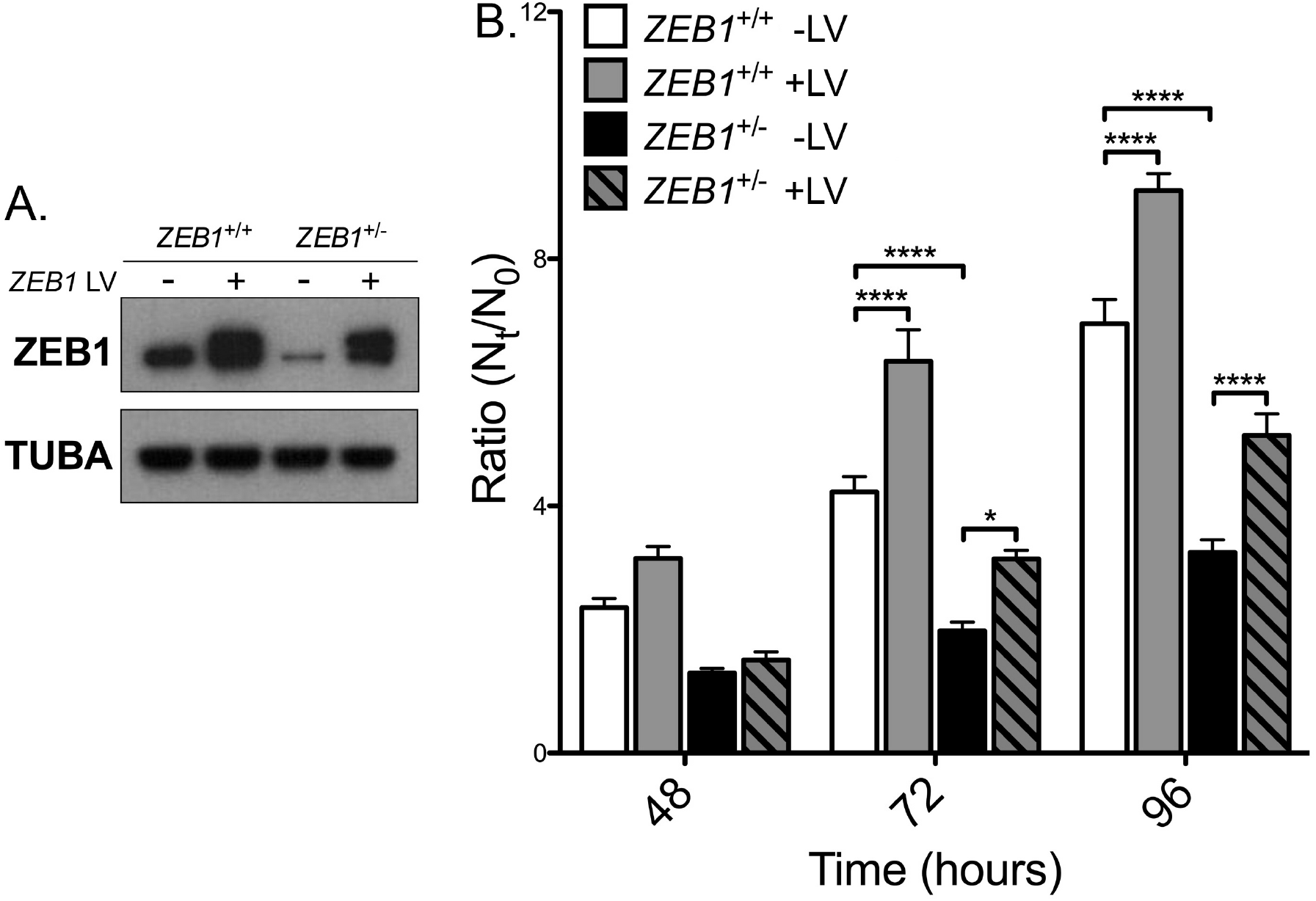
ZEB1 reduction impairs CEnC proliferation capacity. (A) Western blotting results showing ZEB1 levels in each of the CEnC lines following transient ZEB1 overexpression with lentivirus (5 days post-transduction). Alpha-tubulin (TUBA) was used as a loading control. (B) Bar graph showing cell proliferation graphed as the ratio of cell number at time t (N_t_) over cell number at 3 hours (N_0_), N_t_/N_0_. Ratios were calculated at 48, 72 and 96 hours post-seeding. Data were represented as the mean ±SEM (n=6). Comparisons were performed using two-way ANOVA (genotype and time) with post-hoc Bonferroni test. *, P<0.05; ****, P<0.0001.

### ZEB1 insufficiency leads to increased CEnC barrier function

To measure the role that ZEB1 plays in CEnC barrier function, we used electric cell-substrate impedance sensing (ECIS). Barrier function was monitored for 96 hours after initial seeding of cells at 100% confluence (Fig 5). *ZEB1*^+/-^ -LV CEnC demonstrated significantly increased impedance (i.e., increased barrier function), compared with *ZEB1*^+/+^-LV cells (p<0.05) (Fig 5A). *ZEB1* reconstitution in *ZEB1*^+/-^ (*ZEB1*^+/-^ +LV) cells decreased CEnC barrier function to a level that was not significantly different from that in *ZEB1*^+/+^-LV CEnC (p>0.05). Similarly, augmentation of ZEB1 levels in *ZEB1*^+/+^ (*ZEB1*^+/+^+LV) cells resulted in a significant reduction in barrier function compared with *ZEB1*^+/+^-LV CEnC (p<0.05). Both cell-cell (*R*_b_, Fig 5B) and cell-substrate (*α*, Fig 5C) adhesion were contributing factors to overall cell barrier function, demonstrating an inverse relationship compared to ZEB1 levels.

**Fig 5.**
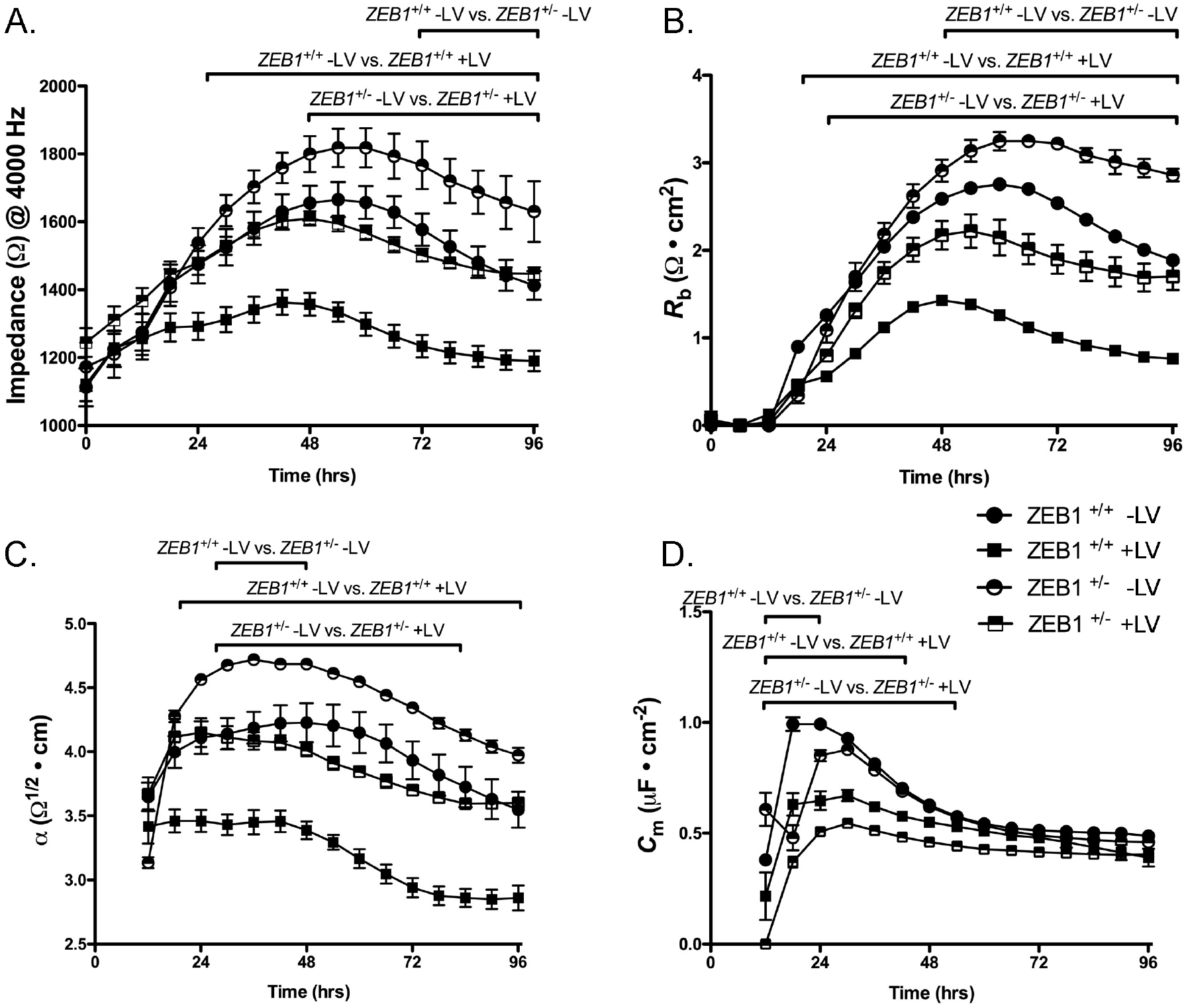
ZEB1 modulates cell barrier function in CEnC. (A) Electrical impedance (Ω at 4000 Hz), a metric of cell barrier function, was measured for up to 96 hours after cells were seeded. (B) Electrical resistance as a result of cell-cell adhesion was modeled from impedance data in (A) and given as *R*_b_ (Ω • cm^2^). (C) Electrical resistance caused by cell-substrate adhesion was modeled from the impedance data in (A) and given as *ケ* (Ω^½^ • cm). (D) Cell membrane capacitance, influenced by membrane complexity and morphology, was modeled from the impedance data in (A) and given as *C*_m_ (μF • cm^-2^). Filled circle: wild type CEnC (*ZEB1*^+/+^ -LV); half-filled circle: *ZEB1* heterozygous CEnC (*ZEB1*^+/-^ -LV); filled square: *ZEB1*^+/+^ cells in which ZEB1 levels were augmented using lentivirus, (*ZEB1*^+/+^ +LV); half-filled square: *ZEB1*^+/-^ CEnC in which ZEB1 levels were reconstituted using lentivirus (*ZEB1*^+/-^ +LV). Data are plotted over 96 hours as the mean ± SEM (n=3,4). Comparisons were performed using two-way (genotype and time) repeated measures ANOVA with post-hoc Bonferroni test. Horizontal bars above curves represent time ranges for the indicated comparisons that demonstrated statistical significance, P<0.05.

### ZEB1 insufficiency does not affect lactate transport in CEnC

Lactate transport is a characteristic function of corneal endothelium, and the original HCEnC-21T line retained this function [24]. Lactate is co-transported across the plasma membrane with protons (H^+^) by lactate monocarboxylate transporters. To determine the effect that ZEB1 levels play in regulating lactate transport, we measured intracellular pH (pH_i_) during various stages of lactate exposure (Fig 6). *ZEB1*^+/+^ -LV cells perfused with lactate buffer demonstrated an influx of H^+^ ions as indicated by the reduction of pH_i_ (Fig 6A). Subsequent perfusion with lactate-free buffer resulted in the efflux of H^+^ and re-establishment of the resting pHi. No significant difference in lactate transport was observed following reduction of ZEB1 (*ZEB1*^+/-^-LV) or with the addition of ZEB1 to either the *ZEB1*^+/+^ or *ZEB1*^+/-^ cells (Fig 6B-D).

**Fig 6.**
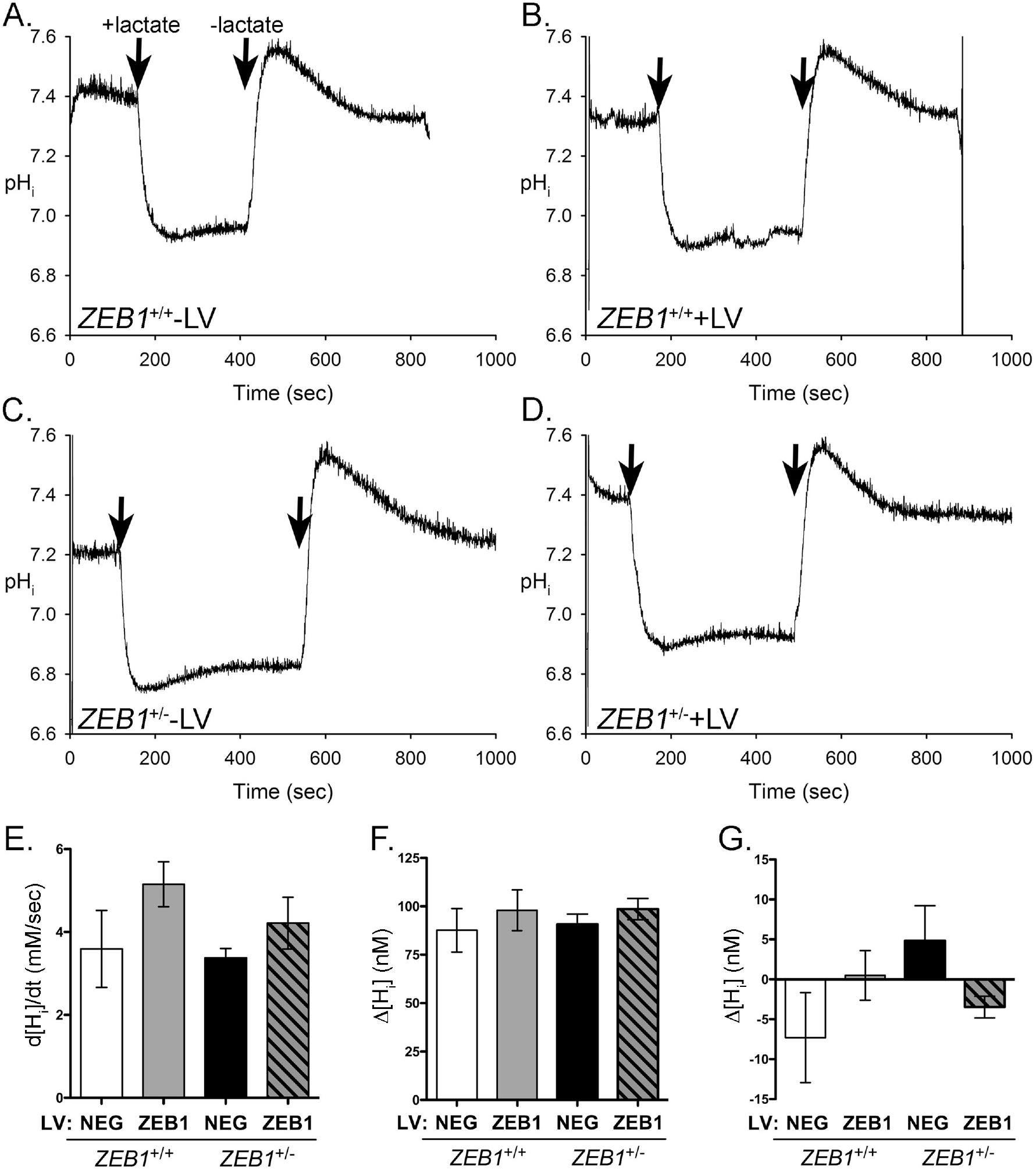
ZEB1 insufficiency does not affect the CEnC response to lactate. (A-D) Traces showing effect of lactate exposure on intracellular pH (pH_i_) in the CEnC lines. Note that lactate is co-transported across the membrane with protons. pH_i_ was calculated from fluorescence measurements of cells pre-loaded with the fluorescent pH indicator BCECF. A resting pHi was established before perfusion with lactate (20mM). Arrows indicate addition or removal of lactate. (E) Bar graph showing the maximum change in intracellular proton concentration ([H_i_], nM) per second (d[H_i_]/dt) after addition of lactate. (F) Bar graph showing the mean of the difference between resting [H_i_] and minimum [H_i_] achieved after addition of lactate. (G) Bar graph representing the mean of the difference between the pre-lactate resting [H_i_] and the post-lactate resting [H_i_]. Data in E-G were represented as the mean ±SEM (n=3). Comparisons in E-G were performed using one-way ANOVA with post-hoc Tukey test. No statistically significant differences were identified.

To assess the dynamics of lactate transport, the change in proton concentration during different phases of lactate perfusion was calculated. After the initial exposure to lactate, a rapid influx of H^+^ ions occurred, leading to a drop in pHi (increase in [H_i_]). The maximum rate of change (d(H_i_)/dt) was calculated and graphed (Fig 6E). While the absolute values (bars) for d([H_i_])/dt appear to be dependent on ZEB1 levels, the association was not statistically significant. We also calculated the difference between resting Hi concentration, achieved before exposure to lactate, and maximum Hi concentration, achieved after the addition of lactate (Δ[H_i_]; Fig 6F). No significant difference was observed in *d*[H_i_ between *ZEB1*^+/+^ or *ZEB1*^+/-^ +/- LV cells. The difference between the pre-lactate resting H_i_ concentration and the post-lactate resting H_i_ concentration was calculated (Δ[H_i_]; Fig 6G). While marked differences in Δ[H_i_] were observed between *ZEB1*^+/+^ and *ZEB1*^+/-^ +/- LV cells, the differences were not statistically significant.

### ZEB1 insufficiency may affect ultraviolet radiation-induced apoptosis in CEnC

Corneal endothelial cell density decreases over an individual’s lifetime, due in part to cell apoptosis. To investigate the effect of reduced ZEB1 on CEnC apoptosis, we exposed *ZEB1*^+/+^ +/- LV and *ZEB1*^+/-^ +/-LV CEnC to ultraviolet C (UVC) for 6 hours and measured phosphorylation of tumor protein 53 (TP53), which is phosphorylated at Serine 15 during apoptosis (Fig 7) [25]. A decrease in phosphorylated TP53 was observed in *ZEB1*^+/-^ -LV cells compared with *ZEB1*^+/+^ -LV cells (Fig 7A). Correspondingly, augmenting ZEB1 levels in both *ZEB1*^+/-^ (*ZEB1*^+/-^ +LV) and *ZEB1*^+/+^ (*ZEB1*^+/+^ +LV) cells resulted in an increase in phosphorylated TP53 compared with *ZEB1*^+/-^ -LV and *ZEB1*^+/+^ -LV cells, respectively. While none of these pairwise comparisons demonstrated statistical significance, it is important to note that the p-value obtained for the 1-way ANOVA was significant (p=0.044), suggesting that the observed means for all four groups, taken together, have a low likelihood of occurring by chance alone (Fig 7B).

**Fig 7.**
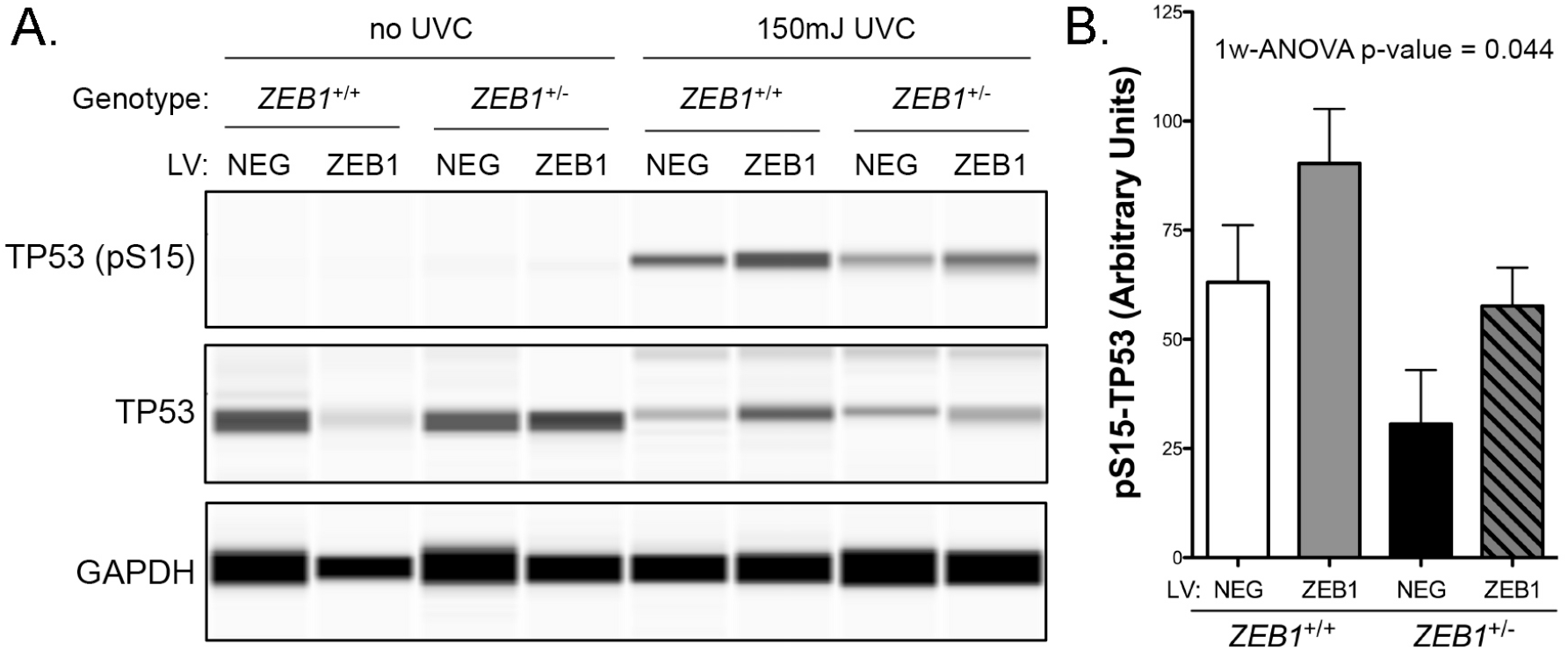
ZEB1 reduction may alter the CEnC response to UVC-induced apoptosis. (A) Western results showing levels of TP53 phosphorylated at Serine 15 in whole-cell lysates prepared from the *ZEB1* CEnC lines treated either with 0 mJ or 150 mJ of UVC. Representative results from three independent experiments are shown. Detection of total TP53 and GAPDH were used as loading controls. (B) Bar graph representing abundance of pS15-TP53 normalized for loading. Data are represented as the mean ± SEM (n=3). Statistical analysis was performed using one-way ANOVA with post-hoc Tukey test.

## DISCUSSION

Cell state transitions are critical processes during embryonic development, tissue remodeling and disease [1]. The transcription factor ZEB1 plays a central role in the regulation of the EMT/MET processes. Changes in ZEB1 expression are sufficient to induce EMT/MET [26], but it is not necessary since other ZEB1-related transcription factors (e.g., SNAIL1, TWIST, OVOL2, GRHL2) have also been shown to mediate these processes [27, 28], with some accomplishing this by directly regulating *ZEB1* transcription [8, 9, 29, 30]. In cancer, the transition from the epithelial to mesenchymal phenotype involves intermediate transition states, which are characterized by the expression of both epithelial- and mesenchymal-associated genes [2–7]. In general, progression towards the mesenchymal state results in a less pronounced epithelial-associated gene expression profile and increased expression of mesenchymal-associated genes.

Contact inhibited and quiescent [31] corneal endothelial cells, when dissociated from the cornea and grown in culture, have demonstrated re-initiation of the cell cycle and transition towards a fibroblast-like (i.e., mesenchymal) phenotype [32, 33], which is associated with an increase in ZEB1 expression [11]. This process is termed endothelial to mesenchymal transition (EnMT) and can be induced by various growth factors and cytokines [34]. As such, there is overwhelming evidence that the terminally differentiated and quiescent corneal endothelial cells retain the potential to undergo a CST towards a fibroblast-like phenotype. Similarly, vascular endothelium has also been observed to undergo EnMT [35], and this is in addition to an EMT-like (epithelial to endothelial) transition that may form the basis for vascular mimicry in cancer [36]. Taken together, these findings raise the possibility that endothelial cells possess the potential to transition to an epithelial-like state, (i.e., endothelial to epithelial transition, EnET).

Evidence for EnET may be found in a disease of the corneal endothelium, posterior polymorphous corneal dystrophy (PPCD). PPCD was first reported in 1916 as a defect of the posterior surface of the cornea [37]. Beginning in the early 1970s, a renewed interest in PPCD culminated in the publication of various comprehensive studies describing clinical [38–42] and histopathologic/molecular features of PPCD [41–45]. Together, these reports provided the first indication that the corneal endothelial cells had gained an epithelial-like phenotype given that the observed morphologic/ultrastructural features and gene expression changes were consistent with such a phenotype. The first transcriptomic study characterizing the gene expression changes in PPCD was reported in 2017, and provided evidence for a widespread increase or ectopic expression of epithelial-associated genes [17]. The genotypes that have been associated with PPCD have also proved to be strong evidence for an MET-like transition of an endothelial to an epithelial phenotype [16]. This is because the genes associated with PPCD have all been reported to play central roles in either EMT (*ZEB1*) [26] and/or MET (*OVOL2* and *GRHL2*) [8, 9, 29, 30].

While the epithelial cell-like features demonstrated by corneal endothelial cells in PPCD have been well characterized, little is known regarding the functional properties of the aberrant endothelial cells. Thus, we recently reported functional consequences of ZEB1 insufficiency in corneal endothelial cells using ZEB1 siRNA [18]. This study was informative, but we recognized its limitations and subsequently developed another cell-based model of PPCD using CRISPR-Cas9 gene-editing technology. We now report the functional impact that stable monoallelic knockout of *ZEB1* has on corneal endothelial cells.

As a prerequisite to performing relevant functional studies, we validated our cell-based model using a transcriptomic approach. Because of the nature of cell culture and immortalization, we expected to observe differences that were not directly relevant to our disease model. Nevertheless, as ZEB1 is robust at mediating EMT in disparate cell types, we concluded that the “background” gene expression was not likely to play a significant role in our study, although this remains a limitation of our model. We demonstrated that the *ZEB1*^+/-^ CEnC possessed a gene expression profile similar to that observed in PPCD, with many epithelial-associated genes demonstrating either increased or ectopic expression in both [17]. Concurrently, we showed that some corneal endothelial associated genes were downregulated in *ZEB1*^+/-^ CEnC, similar to that observed in PPCD. In addition, the observation that reconstitution of the *ZEB1*^+/-^ cells with exogenous ZEB1 caused them to regain a wild type-like (*ZEB1*^+/+^) gene expression profile was particularly notable evidence for the potential utility of gene therapy for PPCD. Taken together, the transcriptomic results indicated that the *ZEB1*^+/-^ cells are an adequate model of PPCD.

The epithelial and mesenchymal (i.e., fibroblastic) cell states can be identified in 2D cultures by characteristic cell shapes associated with each cell state [46]. Epithelial morphology is characterized by a combination of flat, polygonal and cobblestone-like cells, while fibroblast morphology is characterized by a combination of stellate, bipolar and elongated cell shapes. We utilized these differences in the epithelial/fibroblast cell morphology to determine the effects of altered ZEB1 expression on the CEnC state. The observation that a reduction of ZEB1 in CEnC leads to a more robust epithelial morphology provides an in vitro correlate for the in vivo observation that ZEB1 haploinsufficiency leads to an epithelial-like phenotype in PPCD. A logical follow up would be to investigate the potential of *ZEB1*^+/-^ cells to stratify in a 3D culture system, since a stratified organization is also a characteristic feature of the corneal endothelium in PPCD.

Cell migration and cell division are regulated by complex systems involving both mechanical and molecular factors [47]. Nevertheless, robust cell adhesion alone (cell-cell and cell-substrate) explains in large part the reduced migration and cell division observed in epithelial cells, in contrast to fibroblastic cells [19, 48, 49]. As such, cell migration and cell proliferation must first invest a large amount of energy in weakening or breaking cell-cell and/or cell-substrate interactions. In contrast, fibroblastic cells, with notably weaker cell adhesions, possess a higher capacity for migration and cell division. Consistent with these features, *ZEB1*^+/-^ migrated and proliferated less than *ZEB1*^+/+^ cells. While the *ZEB1*^+/+^ cells are not characterized by a fibroblastic phenotype, we postulate that endothelial cells reside in a state between epithelial and mesenchymal (fibroblastic), which is consistent with its hybrid, epithelial/mesenchymal gene expression profile.

The corneal endothelial cell layer is a semipermeable membrane that actively transports substrates in a unidirectional (stroma to aqueous) manner [50, 51]. The endothelium transports water from the corneal stroma to the anterior chamber, thereby maintaining a relatively dehydrated state to achieve/maintain corneal clarity. Because severe cases of PPCD are characterized by endothelial decompensation and edema [12, 52], the net transport function of the endothelium must therefore be impaired. The impairment can occur as a consequence of changes in the expression/targeting/function of membrane solute transporters and/or the physical barrier established by a combination of increased cell-cell, cell-substrate adhesion or stratification of the diseased PPCD epithelial-like cells. In the case of the former, we examined lactate transport, which is a key functional property of the endothelium [53]. We found no significant impact on lactate transport in *ZEB1*^+/-^ cells. While this indicates that ZEB1 insufficiency does not negatively impact lactate transporter function, other transporters may be affected, which warrants further study. An impact on cell adhesion may also have an impact on solute transport as it may establish a significant physical barrier to solute transport. To this end, we observed a significant change in barrier function established by cell-cell and cell-substrate adhesion, with *ZEB1*^+/-^ cells demonstrating significantly greater cell adhesion compared with *ZEB1*^+/+^ cells. This result is consistent with previous reports demonstrating a role for ZEB1 in cellular adhesion [21–23]. Another potential cause of impaired transport function is the establishment of a stratified organization of the epithelial-like cells in PPCD, which would provide an additional physical barrier to the flow of solutes across the corneal endothelium.

Significant cell loss is observed in some PPCD cases, suggesting a potential role for cell death in these cases [12, 52]. In ZEB1 knockdown experiments using siRNA, we demonstrated that reduction of ZEB1 led to an increased sensitivity to UV-induced apoptosis, but not to doxorubicin-induced apoptosis [18]. Herein, in a stable ZEB1 knockdown model, we demonstrate no statistically significant impact of ZEB1 deficiency (or ZEB1 augmentation/rescue) on apoptosis when a pairwise statistical analysis is performed. However, statistical analysis of the collection of means did demonstrate statistical significance. In addition, the positive correlation of ZEB1 with the observed means (i.e., decreased ZEB1 associated with decreased apoptosis and increased ZEB1 associated with increased apoptosis) suggest that ZEB1 may play a role in UVC-induced apoptosis.

In summary, PPCD is characterized by a CST that is consistent with the EMT/MET pathways. This change is marked by gene expression changes consistent with an MET-like transition, and is characterized by the reduction in *CDH2* and an increase in *CDH1* expression, the so-called cadherin switch, a classic feature of EMT/MET. Clinical, histopathologic, genetic and molecular features of PPCD endothelium strongly support a model of disease consistent with a MET-like process (Fig 8). In addition, a majority of the cellular processes investigated in *ZEB1*^+/-^ cells demonstrated results consistent with an epithelial-like phenotype compared with mesenchymal/fibroblastic phenotype. Notably, reconstitution of *ZEB1*^+/-^ cells with exogenous ZEB1 showed the potential clinical utility of *ZEB1* gene therapy with the rescue of the observed epithelial-associated functional phenotypes. Therefore, we propose EnET as a distinct MET-like process important in corneal endothelial biology, with ZEB1 as a key regulator of this transition.

**Fig 8.**
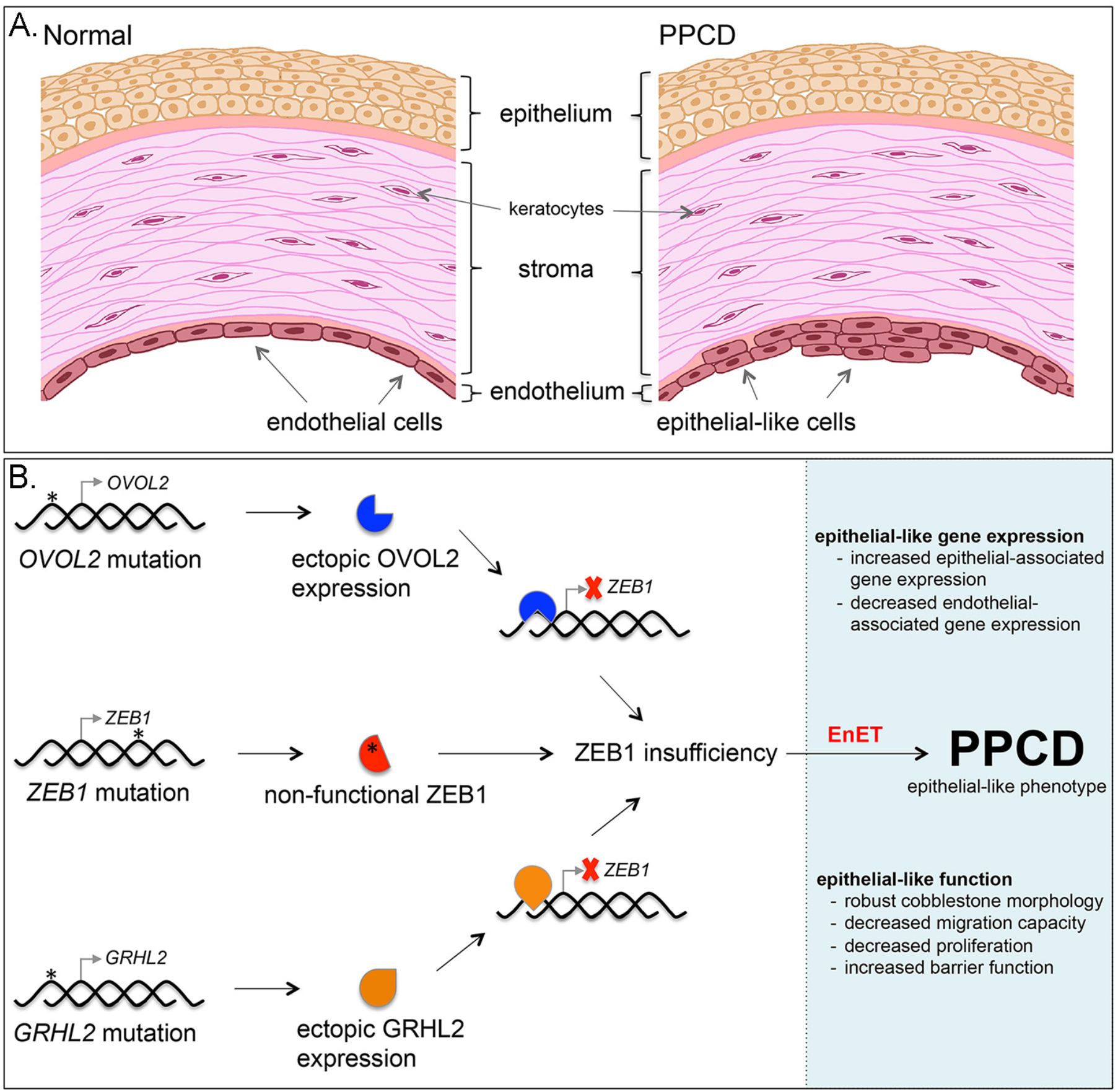
Model for the role of ZEB1 in PPCD characterized by EnET. (A) Illustration of the cornea depicts the three main cellular layers, the anterior stratified organization of the epithelial cells comprising the epithelium, the collagen-rich stroma containing dispersed keratocytes, and the posterior corneal endothelium, which is characterized by a monolayer of corneal endothelial cells. In PPCD, the corneal endothelium is characterized by foci of epithelial-like cells present in a stratified organization, characteristic of the corneal epithelium. (B) Schematic of the genotype-to-phenotype model of PPCD. Truncating mutations (*) in *ZEB1* were the first mutations associated with PPCD. The non-functional mutant protein (red symbol with asterisk) leads to ZEB1 insufficiency and endothelial to epithelial transition (EnET), which forms the basis for the characteristic clinical and histopathologic features of PPCD. Mutations in the promoter region of *OVOL2* or *GRHL2* release intrinsic repression of these genes and lead to ectopic production of their respective transcription factors in the corneal endothelium. OVOL2 (blue symbol) and GRHL2 (orange symbol) are known to directly repress *ZEB1* gene transcription (red X) by binding to the *ZEB1* promoter. Consequently, *ZEB1* transcription is reduced, leading to ZEB1 insufficiency and EnET.

## MATERIALS AND METHODS

Corneal specimens from individuals with posterior polymorphous corneal dystrophy were obtained under a University of California at Los Angeles Institutional Review Board approved protocol (UCLA IRB no. 11–000020). Informed written consent was obtained from all human subjects according to the tenets of the Declaration of Helsinki.

### Cell culture

All CEnC lines in this study were generated from HCEnC-21T cells, an immortalized human corneal endothelial cell line. Cells were maintained and cultured using cell culture-grade plastic flasks coated for 2 hours with a mixture consisting of 40 μg/cm^2^ chondroitin sulfate (Sigma-Aldrich), 40 ng/ cm^2^ laminin (L4544; Sigma-Aldrich), and Dulbecco’s PBS. The cells were grown in a 1:1 mixture of F12-Ham’s medium and M199 medium, supplemented with 5% fetal bovine serum (Atlanta Biologicals), 20 μg/mL human recombinant insulin (Thermo Fisher Scientific), 20 μg/mL ascorbic acid (Sigma-Aldrich), 10 ng/mL recombinant human fibroblast growth factor (basic), 100 μg/mL penicillin (Thermo Fisher Scientific), and 100 μg/mL streptomycin (Thermo Fisher Scientific). HEK293T cells were grown in DMEM supplemented with 10% fetal bovine serum, 100 μg/mL penicillin and μg/mL streptomycin. The cell lines were maintained in a humidified chamber containing 5% CO_2_.

### Cell line authentication

The HCEnC-21T cell line was produced from primary human corneal endothelial cells (sourced from cadaveric corneas) using telomerase immortalization [24]. The authors showed that the cells retained gene expression and functional characteristics of corneal endothelial cells. In addition, after we obtained the cells (a gift from Dr. Ula Jurkunas), we characterized them using a transcriptomics approach and identified the expression of a number of genes distinct for ex vivo human corneal endothelial cells [11]. In this same study, we identified high expression of the human telomerase reverse transcriptase gene, confirming the method used to immortalize the cells. We also performed short tandem repeat (STR) analysis for the *ZEB1*^+/+^ and *ZEB1*^+/-^ cell lines. Genomic DNA was isolated from the cell lines using the FlexiGene DNA Kit (Qiagen). Subsequently, authentication was performed using the PowerPlex 16 System (Promega), a multiplex STR system that complies with ANSI/ATCC ASN-0002-2011 guidelines for cell line authentication. The STR profiles generated for the cell lines were a perfect match to the STR profile of the parental cell line [18].

### Generation of *ZEB1*^+/-^ cell line using CRISPR-Cas9

#### In Silico guide RNA (gRNA) Design

We designed our gRNA (Sigma Aldrich) to target the Cas9 nuclease to exon 4 of *ZEB1* to ensure that the encoded mutant proteins were dysfunctional and that all known splice variants would be affected (S2A Fig.). The design was performed using the crispr.MIT.edu design tool that identifies optimal target sequences with a minimum of potential off-target sites using the hg19 genome build (S2B and S2C Figs).

#### Transfection of HCEnC-21T Cells

The selected gRNA was hybridized to a complementary strand, and was subsequently ligated into pSpCas9(BB)-2A-Puro (PX459) plasmid, a gift from Dr. Feng Zhang (Addgene plasmid #62988) [54] (S2D Fig.). Successfully transfected cells were selected and identified using media containing puromycin. Transfection of the cells was verified by performing Western blots to confirm the presence of the Cas9 protein (S2E Fig.).

#### Clonal Expansion and Characterization

Limiting dilutions were performed to isolate single cells in 96-well plates. Viable cells identified by microscopy were allowed to grow to confluence and were passaged and transferred to 24-well plates. Mutants and controls clones were identified by Sanger sequencing and were transferred to 12-well plates. Genomic DNA was isolated from the clones using QuickExtract (Qiagen) and *ZEB1* was screened in each clone using Sanger sequencing (Laragen). CRISP-ID was used to predict the sequences for each of the two alleles from the Sanger sequencing data that was generated using diploid gDNA template (S3A Fig.) [55]. ZEB1 protein levels were measured using an anti-ZEB1 rabbit monoclonal antibody (AB_1904164) diluted to 1:500 in 0.1% non-fat dried milk in Tris buffered saline solution containing Tween 20.

#### Allele-specific sequencing

Amplicons generated from exon 4 from selected clones were subcloned into plasmid vectors using a TA-cloning kit (Thermo Fisher Scientific)(S3 Fig.). Plasmids containing one of two exon 4 alleles were isolated and screening of the cloned insert was performed using Sanger sequencing (Laragen). The allele-specific sequencing method provided an unambiguous means for identifying indels in each respective allele after CRISPR-Cas9 gene editing. Clones were further characterized by Western blot to detect ZEB1 protein levels and phase-contrast microscopy to assess cell morphology (S3B and S3C Figs). Clones 11 (*ZEB1*^+/+^) and 12 (*ZEB1*^+/-^) were selected to establish cell lines representative of each genotype that was used in this study (S4 Fig.).

#### Screening of off-target sites

As off-target editing by the CRISPR-Cas9 technique may alter cell function in unpredictable ways, identification and screening of potential off-target sites was performed. Primers were purchased from Integrated DNA Technologies and designed to screen each of the top 10 off-target sites predicted by the crispr.MIT.edu tool (S1 Table). Sequencing of each site was performed using Sanger sequencing (S5 Fig.).

### ZEB1 lentivirus production

HEK-293T cells were transfected with a transfer plasmid (pReceiver-Lv215) containing *ZEB1* cDNA of transcript variant 2 (NM_030751.5) (GeneCopoeia) and a 3^rd^-generation packaging system. Cell transfection was performed using LTX transfection reagent with Plus Reagent (Thermo Fisher Scientific) in antibiotic-free medium. Viral supernatants were collected and large particulates were pelleted by centrifugation at 3000 RPM in a swinging bucket rotor (Beckman Coulter). Cleared supernatants were filtered through a 0.45 um syringe filter (Fisher Scientific) and the viral particles were concentrated in an Optima LE8-80K ultracentrifuge (Beckman Coulter) at 25,000 RPM for 90 min at 4°C using the SW28 rotor. Pelleted viral particles were resuspended in 25 uL DPBS for every 10 mL of viral supernatant. Total viral particles were determined by p24 ELISA, performed by the UCLA Integrated Molecular Technologies Core, and the infection units were determined by transduction of HCEnC-21T cells with diluted virus. Subsequently, transduction of the CEnC lines was performed at a multiplicity of infection (MOI) value of 10. Infection was facilitated with the addition of 8 ug/mL of hexadimethrine bromide (Sigma-Aldrich).

### Generation of *ZEB1* transgenic *ZEB1*^+/+^ and *ZEB1*^+/-^ cell lines

While rescue of the ZEB1 insufficiency phenotype was observed for cell proliferation and cell barrier function using transient ZEB1 expression, assay for other cellular functional processes did not demonstrate rescue of the *ZEB1*^+/-^ phenotype following transient reconstitution with ZEB1. To account for the possibility that only a prolonged/constitutive reconstituted expression of ZEB1 was capable of inducing rescue, we generated *ZEB1* transgenic cell lines harboring either the *ZEB1*^+/+^ or *ZEB1*^+/-^ genotype. CEnC were transduced with either empty or ZEB1 lentivirus. Five days after transduction, cell clones were isolated and expanded using the limited dilution method by seeding 0.5 cells/well of a 96-well plate. Several clones were expanded for each CEnC group (*ZEB1*^+/+^ -LV, *ZEB1*^+/+^ +LV, *ZEB1*^+/-^ -LV and *ZEB1*^+/-^ +LV), and were subsequently characterized by cell morphology and ZEB1 Western blot (rabbit monoclonal anti-ZEB1 antibody, AB_1904164). Three clones per cell group were chosen as independent biological lines (12 clones total, 3 per CEnC group). Assays to assess cell migration, cell morphology and lactate transport function were performed with each of the respective clones representing a single independent sample (n=1), so that three independent clones (n=3) were used for statistical analysis. In addition, these 12 clones were used for RNA-seq and qPCR analysis.

### RNA-sequencing and transcriptomic analysis

RNA was isolated from the *ZEB1* CEnC lines and RNA-seq libraries were prepared using the KAPA mRNA HyperPrep Kit using an automated liquid handler (Janus G3 – PerkinElmer) according to manufacturer’s instructions at the UCLA Institute for Quantitative and Computational Biology. Libraries were sequenced on the Illumina HiSeq 4000 platform by the UCLA Broad Stem Cell Research Center High-Throughput Sequencing Core Resource. All RNA-seq data contains single-end 50 base pair reads, which were aligned (grch38.p12) and transcripts quantified (homo sapiens Ensembl Annotation Release 92) using the kallisto (v0.44.0) program [56]. Quantities were given in transcripts per million (TPM). Differential gene expression analysis was performed with the Sleuth (v0.30.0) R-package [57]. Differential expression was tested using a likelihood ratio test, and corrected for multiple testing using the Benjamini-Hochberg correction. The following thresholds defined differential expression: fold change (fc)> 2, TPM>15 and p-value<0.5 (PPCD data); fc>1.5, TPM>0.6 and p-value<0.2 (*ZEB1* CEnC lines). Because the PPCD data was generated from two samples, we used a p-value of 0.5 to reduce false-positives. Given the relatively high variation between the cell lines within each group, a p-value of 0.2 was used for DGE. While this may have increased the number of false-positives, the use of this p-value allowed genes known to be involved in EMT or regulated by ZEB1 to be considered differentially expressed. We generated heatmaps using the *pheatmap* function within the pheatmap (v1.0.10) R-package. RNA-seq data were obtained from the GEO DataSets database (PPCD endothelium, accession number GSE90489; ex vivo endothelium and epithelium, accession GSE121922). RNA-seq data for the cell lines were submitted to GEO DataSets and assigned accession number GSE121680.

Distribution of differentially expressed genes in PPCD endothelium versus evCEnC and evCEpC was statistically analyzed using two methods, a bootstrap approach and the hypergeometric statistical test. The hypergeometric test (hgt) was performed as previously described [58, 59] and was computed in R using the *dhyper* function with significance defined as p<0.05. For the bootstrap method, we combined the Law of Large Numbers and the Central Limit Theorem to create normal sampling distributions centered on the population mean for each of the combinations of gene pools that were examined. We wrote a simulation in R (https://zenodo.org/badge/latestdoi/154679145) and performed 10,000 iterations to create a sampling distribution of the possible outcomes. The number of genes observed by experiment was compared to the sampling distribution to find the probability (p-value) that the number observed by experiment could have occurred by chance. Significance was defined by 0.95 > p < 0.05.

### Immunohistochemistry

Full-thickness PPCD cornea obtained at the time of surgery and cadaveric donor cornea obtained from an eye bank were fixed in 10% Formalin and embedded in paraffin. Tissue was sectioned at a thickness of 5 μm and affixed to a frosted glass slide. Sections were deparaffinized in xylene and rehydrated in an alcohol series. Antigen retrieval was performed with Proteinase K digestion, and tissue was subsequently blocked in 5% goat serum and 1% bovine serum albumin. CLDN1 was detected using a rabbit monoclonal antibody (D5H1D; CST13255), and ADCYAP1R1 was detected using a rabbit polyclonal antibody (AB_777009). Antibodies were diluted 1:1000 in blocking buffer. Detection was performed using an anti-rabbit Alexa fluor conjugated secondary antibody and visualized using a confocal fluorescence microscope.

### Quantitative polymerase chain reaction

Quantitative polymerase chain reaction (qPCR) was performed to validate the level of *ZEB1* gene expression in the *ZEB1* CEnC lines. First-strand synthesis was performed with the SuperScript III First-Strand Synthesis kit (Thermo Fisher Scientific) using oligo-dT primers and 100ng of total RNA. Quantitative PCRs were performed on the LightCycler 480 System (Roche) using the KAPA SYBR FAST qPCR Kit (KAPA Biosystems) and ZEB1-specific oligonucleotide primers (Forward, 5’-TTACACCTTTGCATACAGAACCC-3’; Reverse, 5’-TTTACGATTACACCCAGACTGC-3’; ID: 291575187c2) obtained from the Harvard Primer Bank database [60–62]. Relative gene expression was obtained by comparison to the housekeeping gene *RAB7* and was calculated by the comparative Ct (2^-ΔCt^) method [63]. Transcript quantities were graphed as 2^-ΔCt^.

### Cell morphology analysis using phase contrast microscopy

Images of *ZEB1* transgenic *ZEB1*^+/+^ and *ZEB1*^+/-^ cell cultures at day 1 (sub-confluent) and at day 3 post-seeding (confluent) were acquired using the Leica DMIL LED inverted microscope (Leica Microsystems) and the N PLAN L 20x/0.35 PH1 objective. Image capture was performed with the Leica DFC3000 G monochrome camera controlled with the Leica Application Suite X software (version 3.0.3.16319). Image analysis was performed using ImageJ 1.51h (National Institutes of Health). Regions of interest (ROI) were created along the major axis of cells using the straight-line tool and collected in the ROI manager. After creating an ROI along the major axis of all cells (excluding those cells along the edge of the image) the ROI length in microns was obtained. A total of three fields, each with hundreds of cells for each cell group (*ZEB1*^+/+^ -LV, *ZEB1*^+/+^ +LV, *ZEB1*^+/-^ -LV and *ZEB1*^+/-^ +LV), were assessed.

### Non-wounding cell migration assay

Cell migration was assessed using a non-wounding method. Two-well silicone inserts (ibidi GmbH), each creating a 500um gap, were placed onto cell culture treated plastic. Cells were seeded into each well and allowed to grow to confluence. When cells reached confluence, cell migration was initiated by removal of the silicone inserts. Progression of cell migration was monitored for 24 hours by phase-contrast microscopy using the BZ-X800 microscopy system (Keyence Corporation of America). Image analysis was performed using ImageJ 1.51h software (National Institutes of Health).

### Cell counting proliferation assay

CEnC proliferation was measured for *ZEB1*^+/+^ and *ZEB1*^+/-^ cell lines transduced with ZEB1 lentivirus or an empty lentivirus, which was used as a negative control. Lentivirus (10 MOI) was applied to *ZEB1*^+/+^ and *ZEB1*^+/-^ cell lines and incubated for 5 days, at which point the cells were trypsinized with 0.25% trypsin, counted using a hemacytometer and seeded at 10% confluence on laminin coated plastic. The remaining cells were either used for barrier function analysis or lysed and prepared for Western blotting, which was used to confirm ZEB1 protein levels in each of the four groups. The newly seeded cultures were incubated for 3, 48, 72 and 96 hours. Cells were collected by trypsinization, counted and graphed as a ratio (N_t_/N_0_, where N_0_ equals the number of cells counted at 3 hours and N_t_ equals the number of cells counted at 48, 72 or 96 hours).

### Electric cell-substrate impedance sensing (ECIS) to measure barrier function

A disposable electrode array slide (8W10E+ ECIS, Applied BioPhysics) was stabilized with F99 medium as per manufacturer’s protocol. The array surface was coated with 40 μg/cm^2^ chondroitin sulfate A (Sigma-Aldrich) and 400 ng/cm^2^ laminin (Sigma-Aldrich) in phosphate-buffered saline (PBS) for two hours. Five days after transduction with lentivirus (10 MOI) the cells were reseeded at 100% confluence within respective chambers of the slide array. Cells were incubated in the arrays at room temperature for one hour to facilitate even distribution of cell attachment. After seeding and preliminary cell attachment, arrays were positioned into a 16-well array station and connected to the ECIS Zθ instrument to measure electric impedance (Ω at 4000 Hz) for 4 days. Cell-cell (*R*_b_, Ω • cm^2^) and cell-substrate (*ケ*, Ω^½^ • cm) adhesion along with cell membrane capacitance (*C*_m_, *μ* F • cm^-2^) were modeled from the electric impedance data obtained at 4000 Hz [64].

### CEnC lactate transport function assay

Lactate transport was measured by monitoring free H^+^ concentration (pHi) with a microscope fluorometer [65] using the fluorescence-based (dual-excitation 500 nm and 440 nm) ratiometric pH indicator BCECF (Thermo Fisher Scientific), which was pre-loaded into the cells prior to lactate exposure. BCECF loading was performed in lactate-free solution (20mM Na gluconate, 120mM NaCl, 1mM CaCl_2_, 1mM MgCl_2_, 2.5mM K_2_HPO_4_, 5mM dextrose and 5mM HEPES, pH 7.4), and fluorescence was monitored until a stable pH_i_ was maintained. Subsequently, the lactate-free buffer was replaced by perfusion with lactate-containing solution (20mM Na lactate in place of 20mM Na gluoconate) for about 200 seconds and then switched back to the lactate-free solution.

### UVC-induced CEnC apoptosis assay

The CEnC lines were seeded and allowed to reach confluence prior to irradiation with UVC. The cells were irradiated with 150 mJ m^-2^ of UVC radiation using a Stratagene Stratalinker 1800. Cells were lysed 6 hours post-UVC irradiation. Whole-cell lysates were prepared and processed for protein detection using the Wes separation 12-230 kDa capillary cartridges (Protein Simple). Separation and detection were performed as per the manufacturer’s instructions. Quantification and data analysis were performed using the Compass for SW software (version 3.1.7; build ID: 1205). The phosphorylation of TP53 at Serine 15 was used as measure of apoptosis progression [25]. Total TP53 levels were detected with a rabbit monoclonal antibody (AB_10695803), phosphorylation at Serine 15 of TP53 was detected with a mouse monoclonal antibody (AB_331741), and total TUBA was detected using a mouse monoclonal antibody (AB_1904178). Antibodies were diluted to 1:500 in manufacturer’s blocking buffer.

## ACKNOWLEDGEMENTS

Support provided by National Eye Institute Grants 1R01 EY022082 (A.J.A.), P30 EY000331 (core grant), the Walton Li Chair in Cornea and Uveitis (A.J.A.), the Stotter Revocable Trust (SEI Cornea Division), an unrestricted grant from Research to Prevent Blindness (A.J.A.), National Institute of Diabetes and Digestive and Kidney Diseases 1R01 DK077162 (I.K.), the Allan Smidt Charitable Fund (I.K.), Ralph Block Family Foundation (I.K.), and U.S. Department of Energy Office of Science, Office of Biological and Environmental Research Program DE-FC02-02ER63421 (M.M).

The authors declare no competing financial interests.

## AUTHOR CONTRIBUTIONS

Ricardo F. Frausto and Anthony J. Aldave conceptualized study.

Ricardo F. Frausto, Doug D. Chung and Ira Kurtz designed experiments.

Ricardo F. Frausto, Doug D. Chung, Payton M. Boere, Vinay S. Swamy, Huong N.V. Duong, Liyo Kao, Rustam Azimov, Wenlin Zhang, E. Maryam Hanser and Austin Kassels performed experiments.

Ricardo F. Frausto, Doug D. Chung, Vinay S. Swamy and Liyo Kao performed data analysis.

Ricardo F. Frausto, Vinay S. Swamy, Liam Carrigan and Davey Wong performed statistical analyses.

Marco Morselli prepared RNA-sequencing libraries.

Ricardo F. Frausto and Marina Zakharevich generated the knockout and transgenic cell lines.

Marina Zakharevich prepared artwork in Figure 8.

Ira Kurtz, Matteo Pellegrini and Anthony J. Aldave supervised study.

Ricardo F. Frausto and Anthony J. Aldave wrote original draft of manuscript.

Anthony J. Aldave acquired funding for study.

All authors reviewed and approved the final manuscript.

