## Supplemental Data for "ZEB1 insufficiency causes corneal endothelial cell state transition and altered cellular processing"

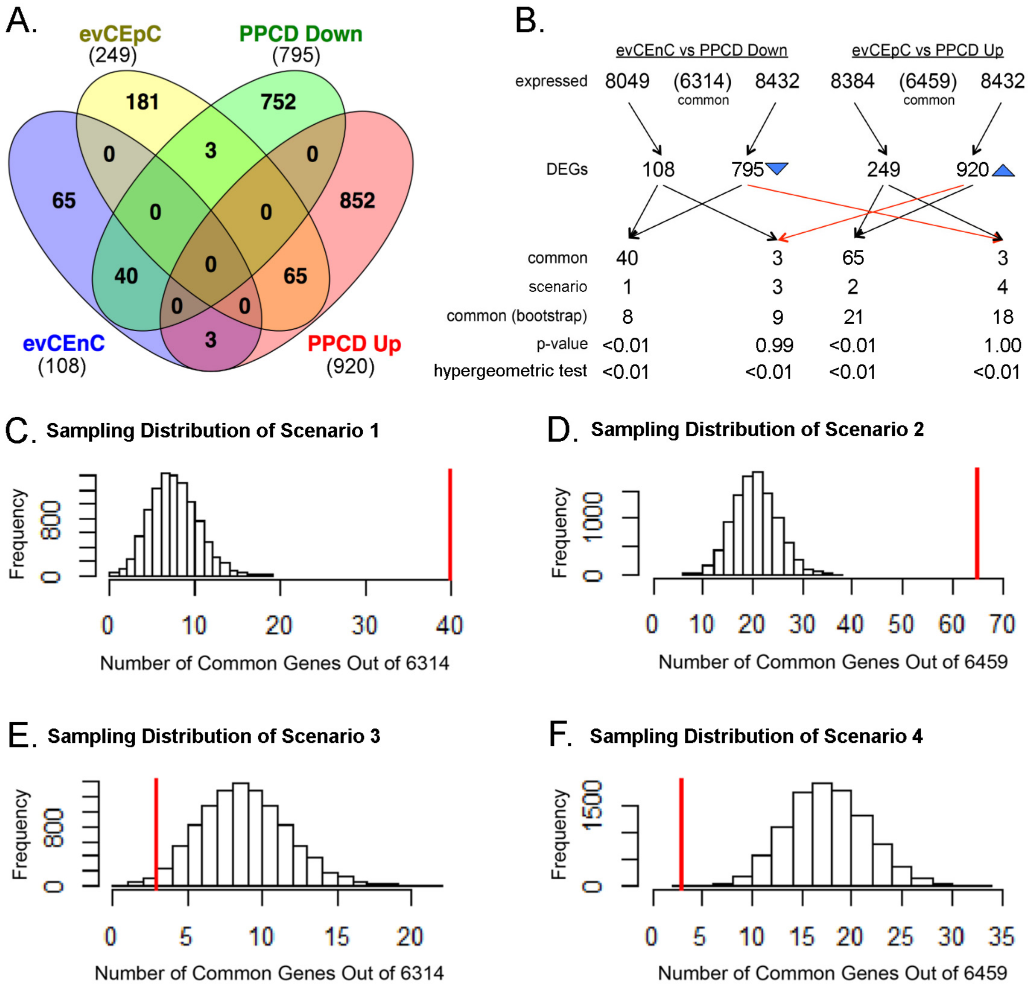


**S1 Fig. PPCD is associated with increased expression of corneal epithelial-specific and decreased expression of corneal endothelial-specific genes.** (A) Venn diagram comparing evCEpC- and evCEnC-specific genes with genes differentially expressed in PPCD. Sixty-eight evCEpC-specific genes were expressed in PPCD endothelium; 65 (96%) demonstrated increased expression. Forty-three evCEnC-specific genes were expressed in PPCD endothelium; 40 (93%) demonstrated decreased expression. (B) Flowchart of number of genes used for statistical testing using a bootstrap approach and summary of results of 10,000 simulations for each scenario. The results of the hypergeometric test (hgt) are also included. Blue arrowheads indicate direction of differential expression. (C) Sampling distribution of scenario 1 where on average 8 genes were expected by chance to be both downregulated in PPCD and evCEnC-specific. Red line indicates observed value (40), which deviates significantly from the mean of the distribution and is not expected by chance alone ($p<<0.01$; hgt $p<0.01$). (D) Sampling distribution of scenario 2 where on average 21 genes were expected by chance to be both upregulated in PPCD and evCEpC-specific. Red line indicates observed value (65), which deviates significantly from the mean of the distribution and is not expected by chance alone ($p<<0.01$; hgt $p<0.01$)). (E) Sampling distribution of scenario 3 where on average 9 genes were expected by chance to be both upregulated in PPCD and evCEnC-specific. Red line indicates observed value (3), which deviates significantly from the mean of the distribution ($p=0.99$; hgt $p=0.01$), and is not expected by chance alone ($p<<0.01$). (F) Sampling distribution of scenario 4 where on average 18 genes were expected by chance to be both downregulated in PPCD and evCEpC-specific. Red line indicates observed value (3), which deviates significantly ($p=1.0$; hgt $p<0.01$) from the mean of the distribution ($p=0.5$), and is not expected by chance alone ($p<<0.01$; hgt $p<0.01$).

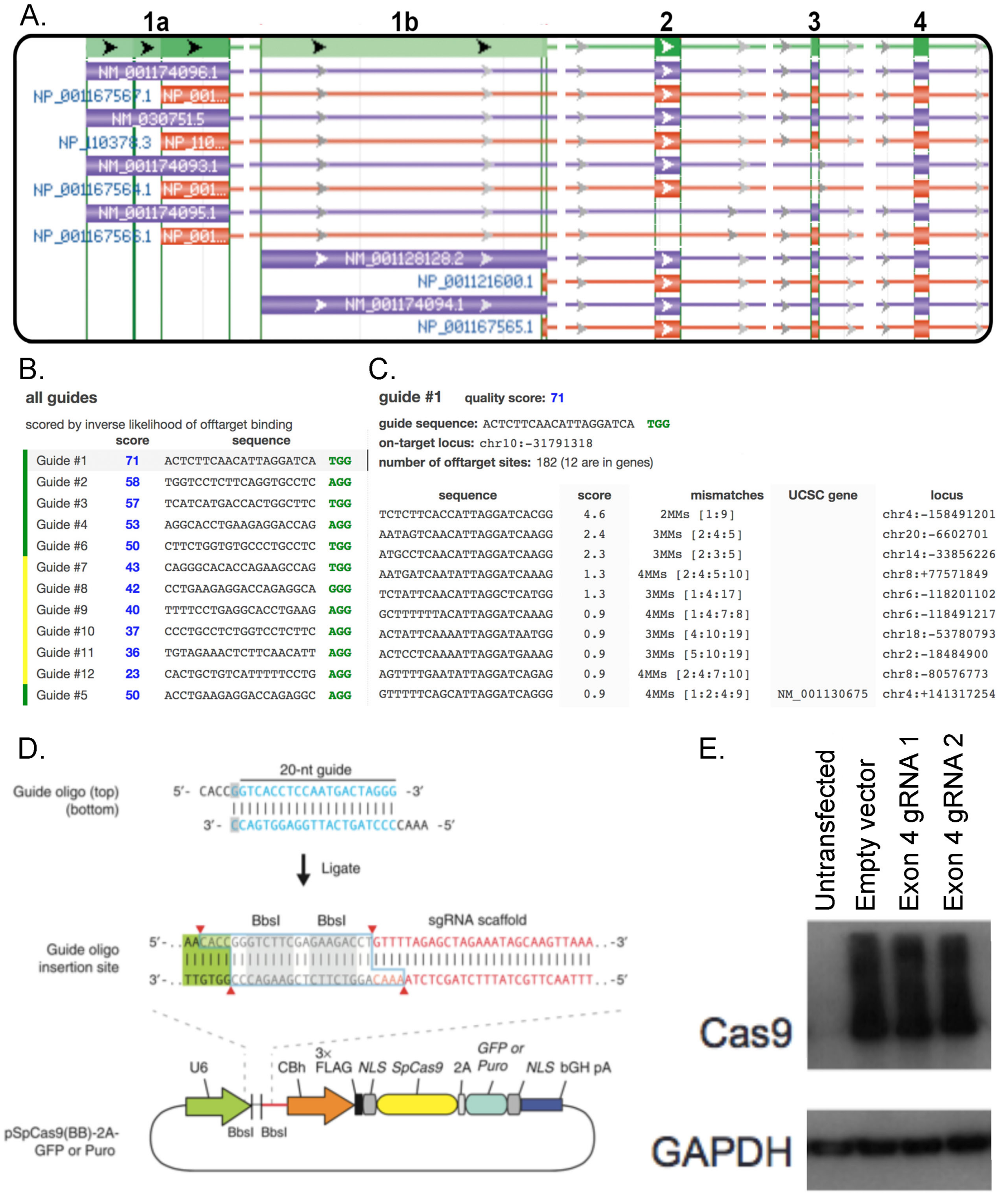


**S2 Fig. Strategy for the generation of the *ZEB1*$^{+/-}$ CEnC line using CRISPR-Cas9.** (A) Screen capture image showing annotated *ZEB1* transcript variants present in the GRCh37.13/hg19 genome build. This build was used because the crispr.MIT.edu guide RNA design tool also utilized the hg19 genome build. Exon 4 was the earliest exon that was present in all *ZEB1* transcript variants and protein isoforms. Exons are indicated by broad colored lines, which are joined by intronic sequences indicated by thin colored lines. Image was modified to accommodate presentation in this figure. Gaps in lines represent intronic sequence that was removed. Exons 5-9 are not shown. (B) List of guides designed to target exon 4 in *ZEB1*. Guides were ranked by score (blue font), which accounts for both on-target and off-target activity. The guide with the highest score (Guide #1) was used for CRISPR-Cas9-mediated editing of exon 4 in ZEB1. The PAM sequence is indicated by green font. (C) The top ten potential off-target sites for guide #1. Several parameters are accounted for in scoring off-target sites, and include number of mismatches, mismatch position and mean pairwise distance between mismatches. Sanger sequencing was used to screen these potential off-target sites (see S5 Fig.). (D) Subcloning strategy for insertion of a guide sequence into pSpCas9(BB)-2A-Puro plasmid. Adapted by permission from Macmillian Publishers: Nature Protocols, Ran FA, et al. 2013; 8(11): 2281-2308, copyright 2013. (E) Western blot demonstrating the presence of Cas9 protein in CEnC whole-cell lysates after transfection with gRNA/Cas9 DNA construct. GAPDH was used as a loading control.

A.

| Clone # | CRISP-ID (allele 1/allele 2) | Allele Sequencing | Predicted Genotype Genetic | Predicted Genotype Protein |
|---|---|---|---|---|
| 9 | insT/delA | insT/**delAinsCT** | -/- | -/- |
| 10 | delGA/insT | delGA/insT | -/- | -/- |
| 11 | WT/WT | WT/WT | +/+ | +/+ |
| 12 | WT/delTC | WT/delTC | +/- | +/- |
| 15 | delTCCTAATGT/delTCCTAATGT | delTCCTAATGT/delTCCTAATGT | -/- | +/- |
| 18 | insT/delA | insT/delA | -/- | -/- |
| 20 | delAAAACCATGA/delAAAACCATGA | delAAAACCATGA/delAAAACCATGA | -/- | -/- |
| 41 | insT/insT | insT/insT | -/- | +/- |
| 44 | insT/delTCCT | insT/delTCCT | -/- | +/- |
| 45 | delCC/delCC | delCC/delCC | -/- | -/- |
| 51 | WT/WT | WT/WT | +/+ | +/+ |

B.

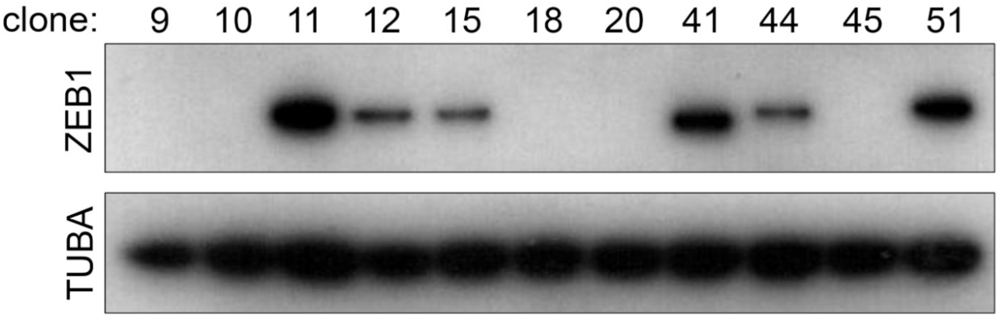


C.

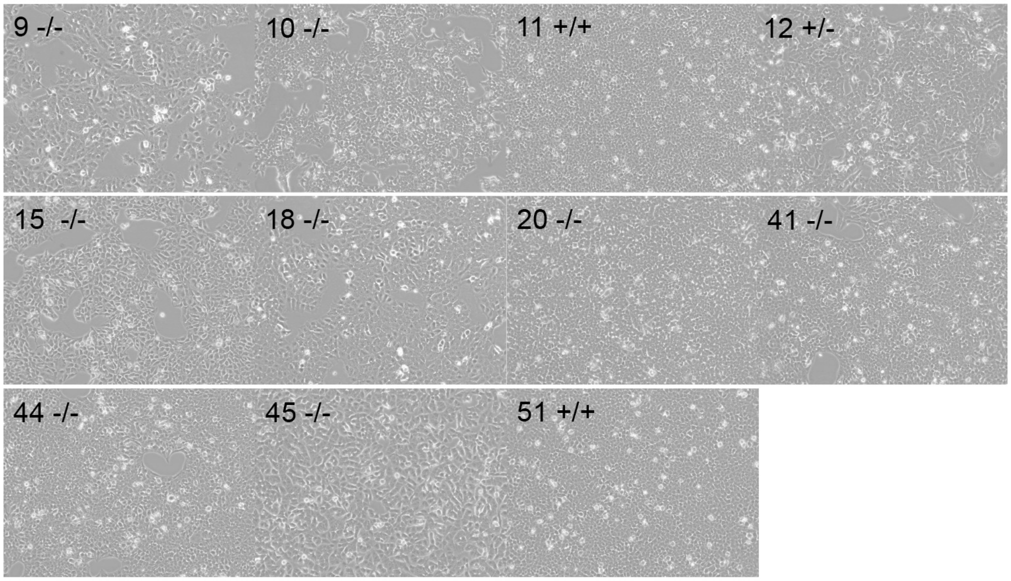


**S3 Fig. Characterization of the CEnC clone lines generated by CRISPR-Cas9-mediated editing of *ZEB1*.** (A) List of selected clones on which additional genetic and molecular characterization was performed. Sanger sequencing of exon 4 of each of the selected clones was performed and the sequencing traces were analyzed using CRISP-ID, which predicted the indels that were introduced after Cas9-mediated DNA cleavage and NHEJ repair. Allele-specific sequencing was performed to validate the indels predicted by CRISP-ID. A simplified description, using "-" for indel and "+" for wild type, of the predicted genotype was compiled for each the DNA sequencing results. (B) Western blotting for ZEB1 shows ZEB1 protein levels in each of the clones. The predicted ZEB1 genotype as interpreted from the Western blot results are show in (A), with clones that did not show consistent predictions shaded in gray. Alpha-tubulin (TUBA) was used as a loading control. (C) Phase-contrast microscopy images showing cultures of each of the cell clones.

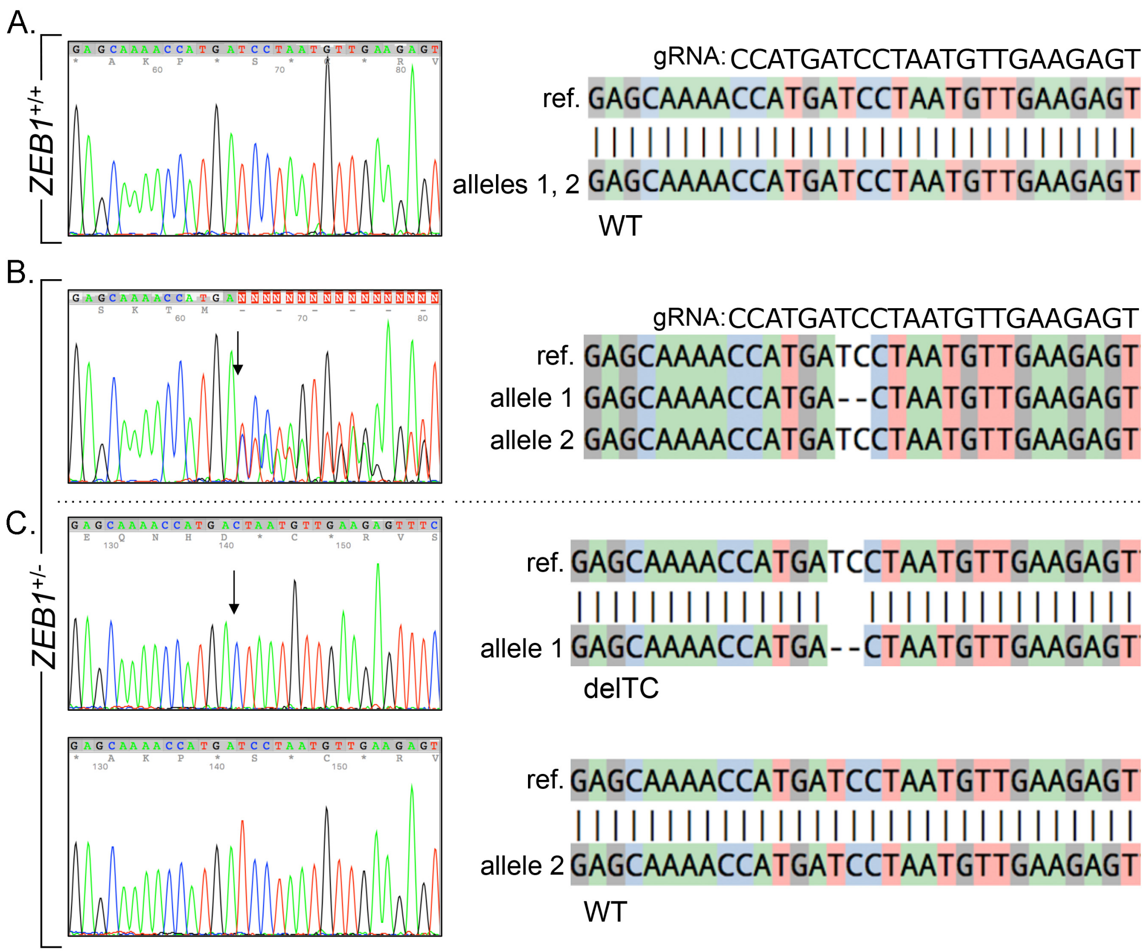


**S4 Fig. Genetic characterization of *ZEB1*$^{+/+}$ and *ZEB1*$^{+/-}$ CEnC lines.** (A) Chromatogram shows Sanger sequencing results of *ZEB1* exon 4 for the *ZEB1*$^{+/+}$ CEnC line. Genomic DNA (diploid) template was used for sequencing. Sequence alignment using CRISP-ID was performed against a reference. Guide RNA sequence is shown above reference sequence. (B) Chromatogram shows sequencing results of *ZEB1* exon 4 for the *ZEB1*$^{+/-}$ CEnC line. Genomic DNA (diploid) template was used for sequencing. Arrow indicates position of the introduction of an indel(s) by NHEJ repair. Sequence alignment to a reference sequence and allele prediction using CRISP-ID shows a mutant allele (delTC) and one wild type allele. (C) Independent sequencing of the individual alleles confirmed that the mutant allele harbors a deletion (delTC), while the second allele was wild type. Arrow indicates position of delTC.

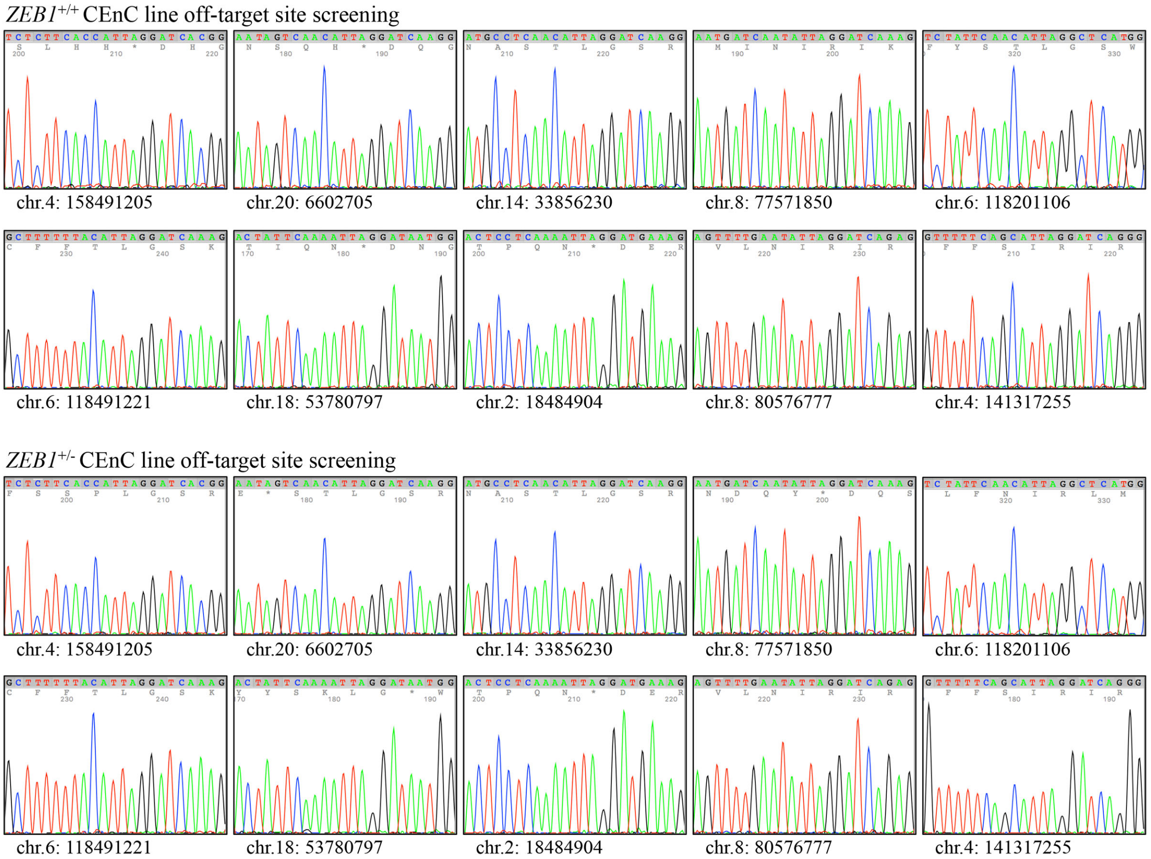


**S5 Fig. Sequencing of potential off-target sites in *ZEB1*$^{+/+}$ and *ZEB1*$^{+/-}$ CEnC lines.** Chromatograms show sequencing results of the *ZEB1*$^{+/+}$ (top set) and *ZEB1*$^{+/-}$ (bottom set) CEnC lines for the 10 off-target sites with the highest scores (see S2C Fig.). Chromosome and position are listed under each chromatogram. Primers for off-target sequencing are in S1 Table.

Table S1. PCR primers for sequencing CRISPR-Cas9 off-target sites.

| **Primer Sequences (5'-3')** | **Amplicon Size (bp)** |
|---|---|
| F1-TGCTAGGATGCCACTAAGCTGT<br>R1-TGGGAGCAGCTCACTTCTCT | 438 |
| F2-ATGGGTTAGAGCAACCAGTGAATTA<br>R2-ACTGTAACCCTCTACTTCTGTAGGC | 401 |
| F3-ATGAGAATCAGGTGGGCGTCT<br>R3-CTGAGGGGCTGACAACACTGA | 438 |
| F4-TACAACCACTGAACCAGACCCTA<br>R4-CACTGACCCCAATGCTTCCA | 402 |
| F5-GTCTTCAGTTCTCTCTCTGAAGCA<br>R5-CGTTGGTCCATGAGCCAAGATG | 419 |
| F6-CTGTGCTCTATTCTGTGAGCCAAA<br>R6-TGTGCCTTAGAAGCAGTCCAACA | 464 |
| F7-AGGCTCATTTGGCGTGCTTT<br>R7-GCTCCCAGCCCTTTGTCCA | 407 |
| F8-GAGCAAAGGCCTTGTCCTATTCA<br>R8-ACATGCAAAAGTGTGCTCCCAA | 401 |
| F9-TGTCTGGTCCTTTGCGTACCAT<br>R9-AAGGGCCTTGACAAACAGGTCA | 416 |
| F10-CACACCCAATCCGACATGCTG<br>R10-GGCACAAACAAAAGGGAGGGAAA | 411 |

Note: annealing temperature used for PCR was 61°C for all primer pairs.
